# Diverse viral *cas* genes antagonize CRISPR immunity

**DOI:** 10.1101/2023.06.24.545427

**Authors:** Mark A Katz, Edith M Sawyer, Albina Kozlova, Madison C Williams, Shally R Margolis, Luke Oriolt, Matthew Johnson, Joseph Bondy-Denomy, Alexander J Meeske

## Abstract

Prokaryotic CRISPR-Cas immunity is subverted via anti-CRISPRs (Acrs), small proteins that inhibit Cas protein activities when expressed during the phage lytic cycle or from resident prophages or plasmids. CRISPR-Cas defenses are classified into 6 types and 33 subtypes, which employ a diverse suite of Cas effectors and differ in their mechanisms of interference. As Acrs often work via binding to a cognate Cas protein, inhibition is almost always limited to a single CRISPR type. Furthermore, while *acr* genes are frequently organized together in phage-associated gene clusters, how such inhibitors initially evolve has remained unclear. Here we have investigated the Acr content and inhibition specificity of a collection of *Listeria* isolates, which naturally harbor four diverse CRISPR-Cas systems (types I-B, II-A, II-C, and VI-A). We observed widespread antagonism of CRISPR, which we traced to 12 novel and 4 known Acr gene families encoded on endogenous mobile genetic elements. Among these were two Acrs that possess sequence homology to type I-B Cas proteins and assemble into a defective interference complex. Surprisingly, an additional type I-B Cas homolog did not affect type I immunity, but instead inhibited the RNA-targeting type VI CRISPR system through sequestration of crRNA. By probing the IMGVR database of viral genomes, we detected abundant orphan *cas* genes located within putative anti-defense gene clusters. We experimentally verified the Acr activity of one viral *cas* gene, a particularly broad-spectrum *cas3* homolog that inhibits type I-B, II-A, and VI-A CRISPR immunity. Our observations provide direct evidence of Acr evolution via *cas* gene co-option, and new genes with potential for broad-spectrum control of genome editing technologies.

## MAIN

Prokaryotic CRISPR-Cas systems use RNA-guided Cas nucleases to provide their hosts with sequence-specific immunity against foreign genetic elements, such as bacteriophages and plasmids^1, 2^. Small fragments of foreign DNA are captured and integrated as “spacer” sequences in the CRISPR locus, which is then transcribed and processed into mature crRNAs^3, 4^. These RNAs guide Cas nucleases in recognition and cleavage of matching targets in foreign nucleic acids^1, 2, 5^. In response to the strong selective pressure imposed by CRISPR immunity, phages and other mobile genetic elements have evolved anti-CRISPR proteins (Acrs), which antagonize the immune effector activities of Cas proteins, removing the barrier to infection^6^. Acrs work via diverse mechanisms to inhibit critical steps of CRISPR immunity, including *cas* gene expression^7, 8^, assembly of CRISPR ribonucleoprotein complexes^9, 10^, recognition of target nucleic acids^11, 12^, and recruitment of effector nucleases^11^. CRISPR-Cas systems are highly diverse immune modules that differ in their *cas* gene sequences, organization, and mechanism of target interference^13^. Most characterized Acrs act via a protein-protein interaction with their cognate Cas protein, and therefore inhibition specificity is almost always limited to a single CRISPR subtype.

How *acr* genes arise within phage genomes is not well understood. While some Acrs have enzymatic activity and likely evolved from enzymes sharing the same fold, a lack of detectable protein homology for most Acrs limits our ability to understand their origins^10, 14–16^. One Acr (AcrIF3) has been shown to mimic the structure of the Cas protein Cas8f to block recruitment of the type I-F CRISPR nuclease Cas2-3^17, 18^. AcrIF3 does not bear significant sequence homology to Cas8f, therefore it is unknown whether this is a case of convergent evolution, or if the two proteins share a common ancestor but have diverged to the point of unrecognizable similarity. Many archaeal viruses encode homologs of Cas4, which normally plays a role in processing newly acquired spacers^19–21^. Some experimental evidence suggests viral Cas4 proteins inhibit spacer acquisition, suggesting that *cas* genes might be co-opted by viruses for CRISPR antagonism^22^. While viral CRISPR-Cas systems are diverse and abundant^23^, no viral *cas* gene has been shown to inhibit the interference stage of immunity, and the extent of *acr* evolution from *cas* genes has not been explored.

*Listeria spp*. have evolved a diverse suite of immune defenses, including four types of CRISPR-Cas systems, to defend against abundant invading phages and mobile genetic elements^24^. The foodborne pathogen *Listeria monocytogenes* is a target of phage-mediated biocontrol efforts, and understanding the anti-defense arsenal of *Listeria* phages holds the potential to enhance this approach^25^. Previous studies have uncovered Acrs encoded by *Listeria* phages, including six that inhibit type II-A and one that inhibits the type VI-A CRISPR-Cas system^9, 12, 26^. Genes encoding these inhibitors are often clustered together in operons, located downstream of phage lysin genes, or within plasmids. Here we investigated the frequency of endogenous Acr-mediated inhibition by screening the functionality of 4 CRISPR types across 62 strains of *Listeria seeligeri.* We bioinformatically predicted *acr* gene candidates and tested them, guided by the results of our functional screen. These efforts uncovered 12 novel *acr* gene families (7 type I-B, 3 type II-C, 2 type VI-A). We found that 3 of these genes bear sequence similarity to type I-B *cas* genes. Two of them (AcrIB3 and AcrIB4) inhibit type I-B by assembling into a defective interference complex. The other (AcrVIA2) is a Cas3 homolog that inhibits the loading, processing, or stability of Cas13-associated crRNAs. Finally, we performed a bioinformatic search for orphan *cas* genes in viral genomes, revealing 358 putative anti-defense loci anchored by a diverse set of type I, II, III, IV, and VI *cas* genes. We experimentally verified one of them in *Listeria,* a Cas3 homolog exhibiting particularly broad-spectrum inhibitory activity against type I-B, VI-A, and II-A CRISPR immunity. Overall, our results exemplify the complex phage-bacteria arms race, and support a mechanism for frequent *acr* gene evolution from *cas* genes.

### Variation in *Listeria seeligeri* genomes affects CRISPR-Cas function

CRISPR-Cas loci can be readily identified by analysis of microbial genome sequences. However, whether these systems provide functional immunity cannot be inferred from sequence analysis alone. We previously established *Listeria seeligeri* as a tractable model for studying type VI-A CRISPR-Cas immunity, and found that *L. seeligeri* strains are also richly populated with type I-B, II-A, II-C CRISPR systems, along with many prophages and plasmids^12, 24, 27, 28^. We sought to determine the extent to which resident mobile genetic elements and prophages affect the function of all four *Listeria* CRISPR-Cas types. We cloned each type into the site-specific integrating vector pPL2e^29^ under the control of a constitutive promoter, and equipped each with a spacer recognizing a target plasmid (Fig. 1A). We first introduced each of these constructs into *L. seeligeri* strain LS1 and confirmed that all four were capable of mediating sequence-specific interference against a target plasmid that was introduced by conjugation (Fig 1B). Next, we integrated each plasmid-targeting CRISPR-Cas construct into 54 out of the 62 *L. seeligeri* strains in our laboratory’s collection, then challenged each of the 216 resultant strains with a cognate target plasmid (Fig. 1C-D and Figs. S1-4). While each CRISPR type remained functional in some of the recipient strains, we observed frequent loss of CRISPR function among the different genetic backgrounds. The loss of CRISPR function we observed for each type did not correlate with the natural occurrence of that type in the tested strains (Fig 1E). We observed either a partial or complete loss of CRISPR-Cas system function in 29% of strains assessed for type VI-A activity, 77% tested for type I-B activity, 36% tested for type II-A activity, and 39% tested for type II-C activity. In some cases, we were unable to determine whether a particular CRISPR type was inhibited, due to low conjugation efficiency of both the target plasmid and a non-targeted control (Fig. 1B, gray bars). The strains tested were least likely to support the function of type I-B, the most common *L. seeligeri* CRISPR type, while most supported the function of type VI-A, which is less abundant. In contrast, type II-C is the rarest type among L. seeligeri strains, yet it frequently lost function when integrated into our tested strains. Collectively, our findings indicate that variation in genetic background affects the function of all four CRISPR types found in *Listeria spp*.

**Figure 1.**
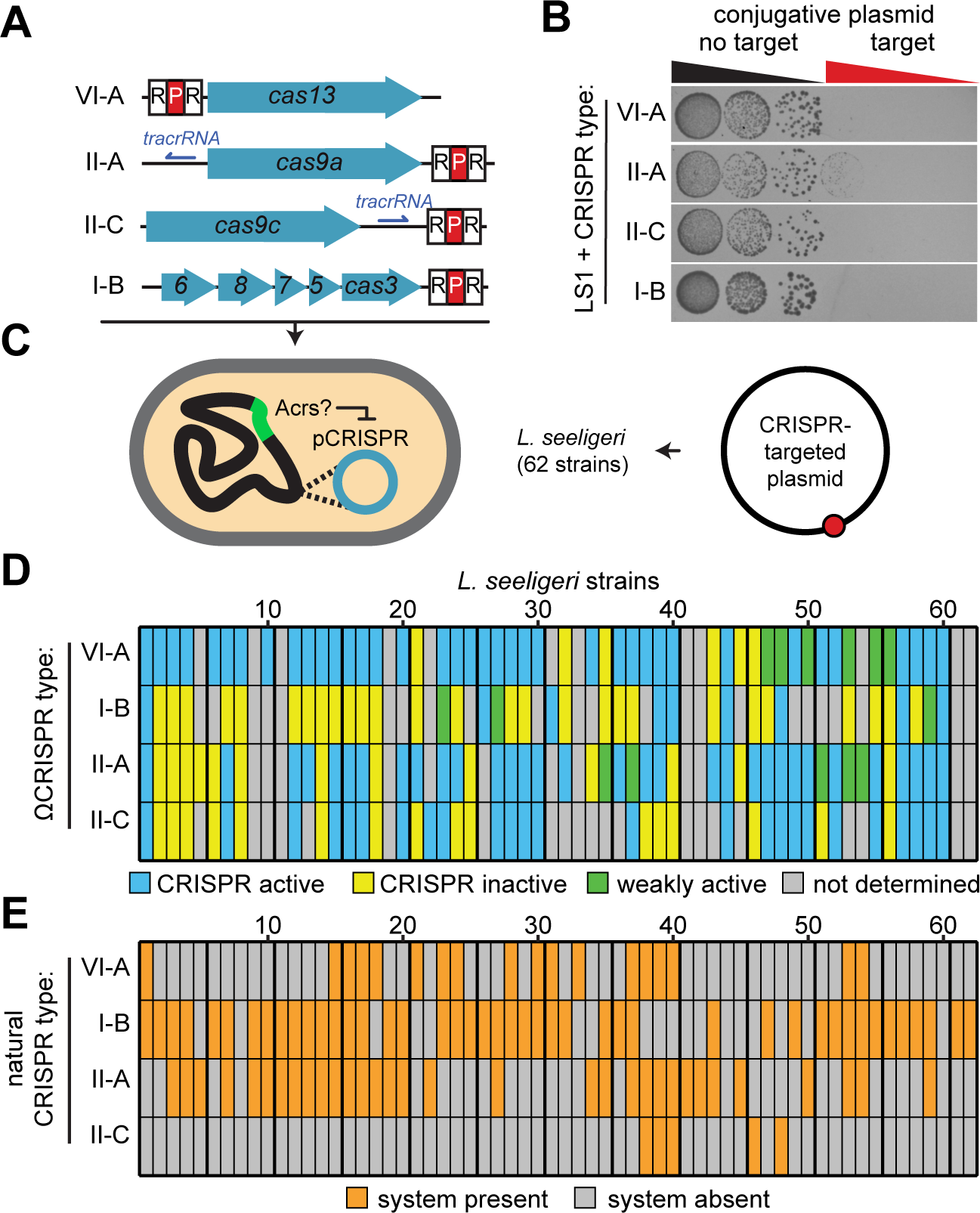
Variation in L. seeligeri genomes affects CRISPR-Cas function. **(A)** Schematic of mobilizable chromosomally-integrating CRISPR-Cas loci each equipped with a single plasmid-targeting spacer. **(B)** Plasmid targeting assay demonstrating sequence-specific interference by all four CRISPR-Cas types in strain LS1. **(C)** Schematic of strategy to detect activity of endogenous Acrs by introducing CRISPR-Cas loci into diverse strain backgrounds and challenging them with target plasmid. **(D)** Functionality of 4 CRISPR types across 62 L. seeligeri strains. **(E)** Natural occurrence of CRISPR types across the strain collection.

### Identification of type I-B, II-A, II-C, and VI-A anti-CRISPR proteins

While the results above could be explained by the variable presence of unknown host factors required for CRISPR-Cas function, we hypothesized that the four CRISPR types might also be inhibited by anti-CRISPR proteins endogenously expressed by the strains in our collection. To identify such inhibitors, we took an iterative guilt-by-association bioinformatic approach that was guided by the results of our functional screen. Six type II-A inhibitor proteins and one type VI-A inhibitor have been previously identified in *Listeria* phage genomes, and Acr genes are frequently clustered in operons associated with prophages or other mobile genetic elements. Therefore, we tested genes located within predicted Acr clusters for the ability to inhibit CRISPR types that could no longer mediate interference when transplanted into the cluster’s host genome. First, we searched each of the *L. seeligeri* genomes in our collection for genes homologous to 81 known Acrs, which resulted in the identification of 25 predicted Acr loci. (Table S2). We examined the genes predicted to be in the same operon as known Acrs in these loci, generating a list of 33 putative anti-defense candidate gene families. Using these new candidates as queries, we searched the genomes again to find new putative anti-defense loci and anti-defense candidate genes, giving priority to genes located between previously identified candidates. We also identified predicted loci and candidate genes by searching *Listeria* genomes available in the NCBI nr and wgs databases. By exhaustively iterating this process, we expanded our dataset to 55 predicted anti-defense loci and 76 anti-defense candidate gene families residing within the 62 *L. seeligeri* genomes in our collection (Fig. S5, Table S2).

Next, we investigated whether the Acr content of each host strain correlated with loss of function for each transplanted CRISPR type (Figs. S5-9). No known type I-B inhibitors exist in *Listeria*. However, of the 13 strains that did not support type II-A CRISPR function, all encoded at least one previously identified type II-A Acr (Fig. S7). Conversely, only 2 of the 32 strains supporting type II-A function contained a cognate *acr* gene. Furthermore, AcrIIA1 inhibits both type II-A and type II-C immunity^8^, and was present in 10 of the 15 strains lacking type II-C function (Fig. S8). Finally, the only known type VI-A *acr* (*acrVIA1*) in *Listeria* was present in a genome incompatible with type VI-A interference, and was absent from all other genomes (Fig. S9). These data suggest that the loss of CRISPR function observed in our screen can largely be explained by host-encoded Acrs. We identified anti-defense candidate genes specifically present in strains that inhibited types I-B, II-C, and VI-A, and expressed each from a plasmid in strain LS1, which does not harbor any anti-CRISPR genes. We then tested whether each candidate inhibited the matching CRISPR type in our plasmid targeting assay (Fig. 2A). We prioritized testing of candidates that were present in inhibitory strains for a given CRISPR type but absent from strains that tolerated function of that type. We ultimately cloned 43 candidate genes, as well as 7 previously identified *acr* genes, and tested each for inhibition of all four CRISPR types (Fig. 2B-C). Of the tested novel candidates, 7 inhibited type I-B (hereafter *acrIB3-9*), 3 inhibited type II-C (*acrIIC7-9*), and 2 inhibited type VI-A CRISPR immunity (*acrVIA2-3*) (see Table S2 for protein sequences). Each of these Acrs were tested against each CRISPR type, but specifically inhibited only one of the 4 types. We also noted that a *L. seeligeri* homolog of the AcrIIA3 protein tested in our assay was a potent inhibitor of type II-C CRISPR, and did not inhibit type II-A, despite being 94.3% identical to *L. monocytogenes* AcrIIA3. While more than one Acr might be active in a given genome, the previously identified and newly discovered Acrs could collectively account for 68% of the inhibition observed in our functional screen.

**Figure 2.**
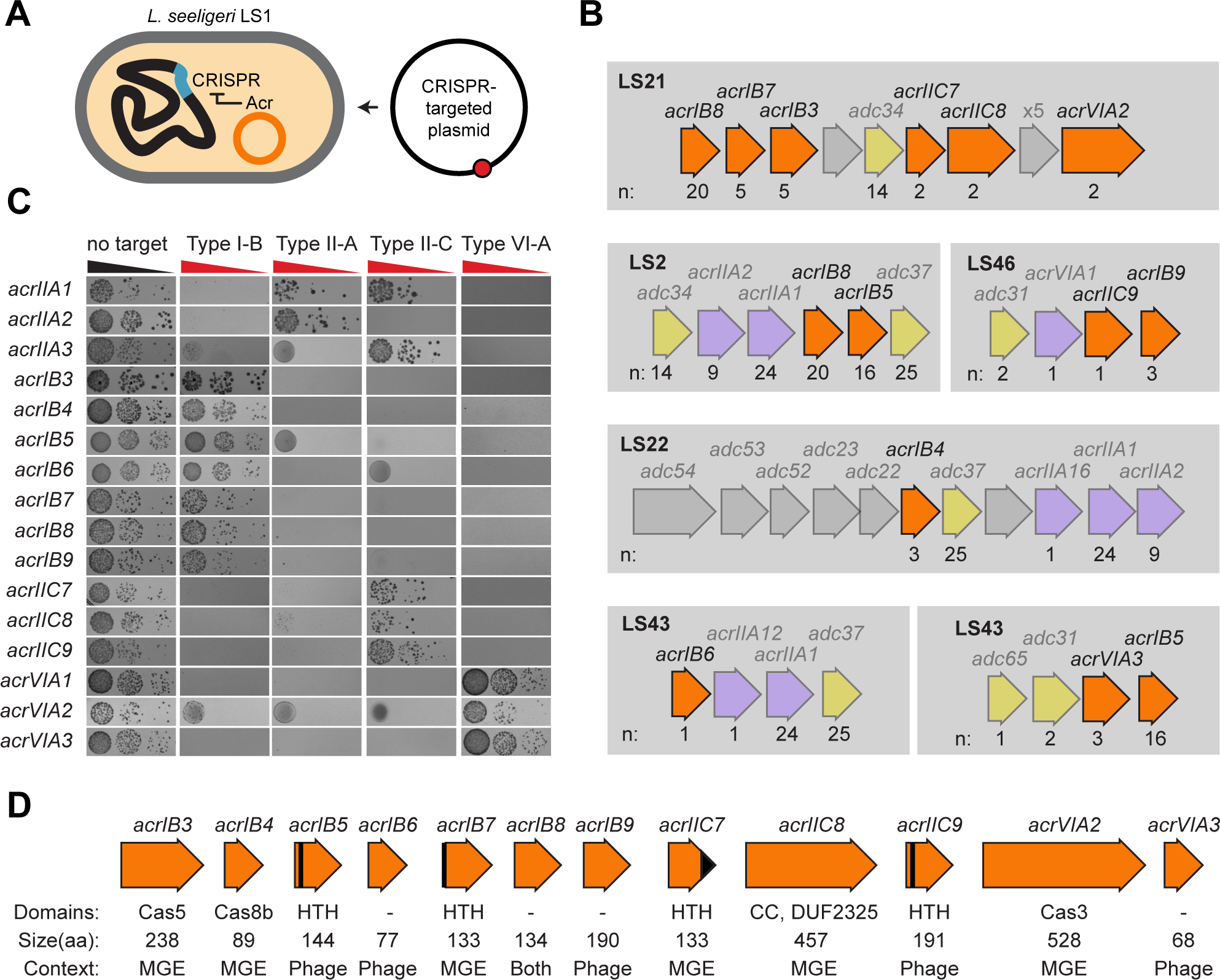
Identification of 12 type I-B, II-C, and VI-A anti-CRISPR families. **(A)** Schematic of strategy to test Acr candidates. Acrs were expressed from a plasmid and introduced into strain LS1 harboring an active CRISPR-Cas system, then challenged with a target plasmid. **(B)** Genetic loci encoding known and novel Acrs. Known Acrs shown in purple. Anti-defense candidate genes used in prediction of Acr loci shown in yellow. Novel Acrs with activity demonstrated here shown in orange. n, number of instances of indicated gene in *L. seeligeri* collection. **(C)** Inhibition spectrum of tested Acrs. Each Acr candidate was tested against all four CRISPR-Cas systems in a plasmid targeting assay. **(D)** Acrs discovered in this study. Predicted protein domains noted, with HTH (helix-turn-helix) domains depicted in black. MGE, mobile genetic element.

In total, we discovered 12 new Acr families, 11 of which each had several homologs present in a variety of *Listeria* species and phages (Fig. 2D, Fig. S10). The occurrence of these *acr* genes was limited to *Listeria*, except for AcrIIC9, which was also found in other *Firmicutes,* notably *Enterococcus.* Genes encoding homologs of AcrIB3, AcrIB4, AcrIB7, AcrIB8, AcrIIC7, AcrIIC8, and AcrVIA2 were found in mobile genetic elements within *Listeria* genomes, while AcrIB5, AcrIB6, AcrIB7, AcrIB9, AcrIIC9, and AcrVIA3 were found in *Listeria* phage genomes. Few of these Acr proteins contained domains of known function, however, we noted that four of them contained HTH domains predicted to mediate DNA binding. Indeed, in addition to its CRISPR inhibition discovered here, we previously demonstrated that the gene encoding AcrIIC9 functions as a negative autoregulator of its own *acr* gene locus^12^. Finally, three of the Acrs shared sequence homology with type I-B Cas proteins, which we discuss in detail below.

### Cascade component homologs inhibit type I-B CRISPR-Cas immunity

Two of the newly discovered type I-B Acr proteins (AcrIB3 and AcrIB4) shared sequence homology with two type I-B Cas proteins (Cas5 and Cas8b, respectively) that assemble into the Cascade complex (Fig. 3A). The AcrIB3 protein shares 38% amino acid identity with the full-length Cas5 protein (Fig. S11A), while the AcrIB4 protein shares 38% amino acid identity with the last 90 residues of the 562aa Cas8b protein (Fig. S11B). We hypothesized that AcrIB3 and AcrIB4 might inhibit type I-B CRISPR immunity by acting as faulty subunits integrated within the Cascade complex. An alternative possibility is that expression of any individual natural Cas protein from a multi-copy plasmid would interfere with Cascade complex assembly via disruption of subunit stoichiometry. To test whether this was the case, we separately expressed AcrIB3, AcrIB4, and their cognate Cas protein homologs Cas5 and Cas8b, and tested their effect on plasmid targeting by the type I-B CRISPR system (Fig. 3B). While the two Acrs potently inhibited interference against the target plasmid, neither *bona fide* Cas protein impacted immunity. We performed BLAST searches of AcrIB3 and AcrIB4 against the NCBI nr database. In addition to numerous true Cas5 and Cas8b protein homologs, we uncovered 45 and 49 unique homologs (respectively) which were not located within CRISPR-Cas loci, and all limited to *Listeria spp*. (Fig. 3C). Our phylogenetic analysis of the proteins uncovered by this search indicated that both Acrs form their own high-confidence clades, suggesting an ancient divergence from their cognate Cas proteins. We therefore conclude that AcrIB3 and AcrIB4 are Cas protein homologs that function as inhibitors of the type I-B CRISPR-Cas system.

**Figure 3.**
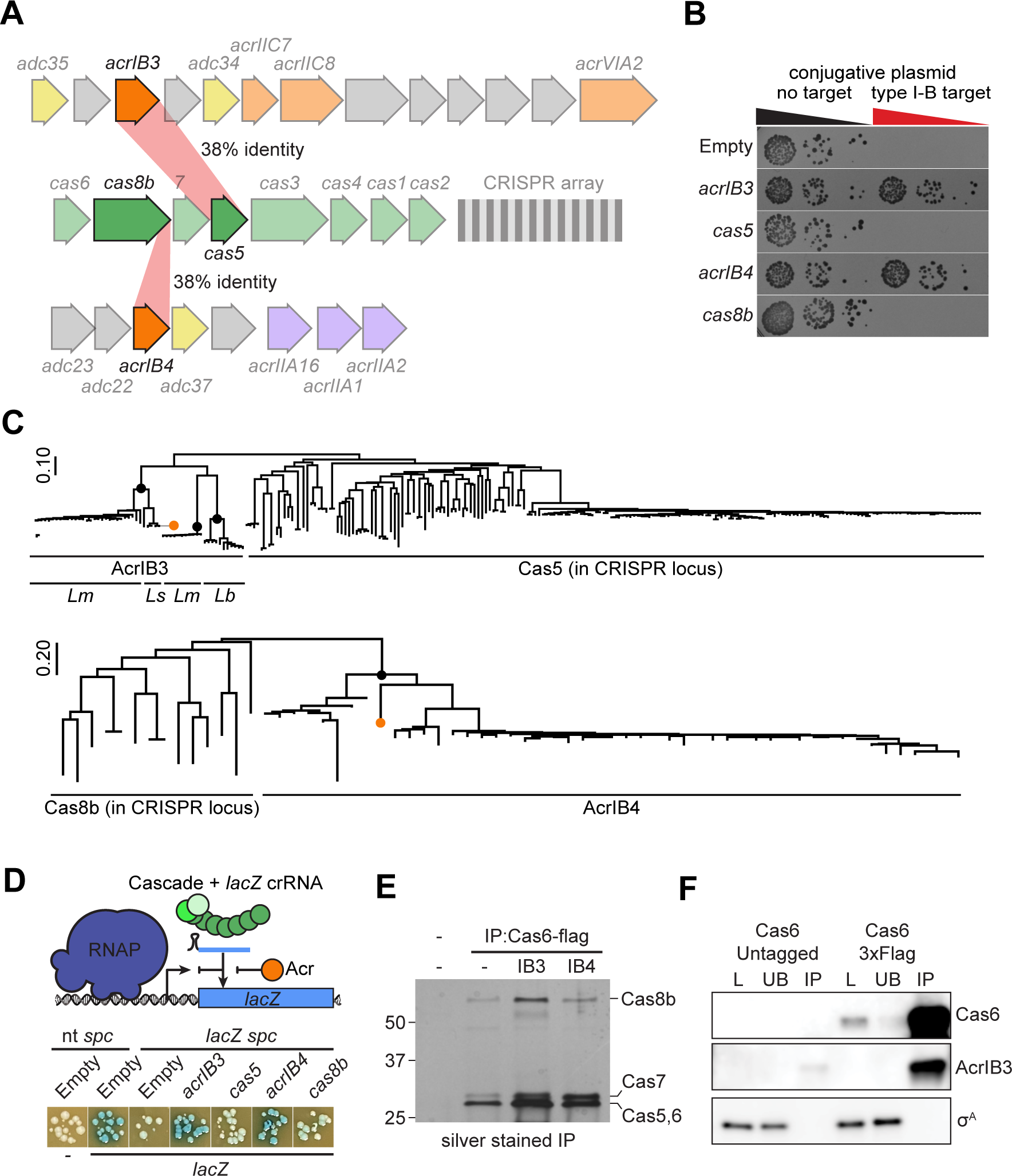
AcrIB3 and AcrIB4 are Cas protein homologs that inhibit type I-B CRISPR immunity. **(A)** Schematic of genetic loci encoding AcrIB3 and AcrIB4 (opaque orange) and type I-B CRISPR-Cas locus. Percent sequence identity between AcrIB3 and Cas5, and AcrIB4 and the CTD of Cas8b are noted. **(B)** Plasmid targeting assay demonstrating that expression of AcrIB3 and AcrIB4, but not their cognate Cas proteins Cas5 and Cas8b, inhibits type I-B CRISPR immunity. **(C)** Predicted phylogeny of AcrIB3 and AcrIB4 homologs uncovered by BLAST search. Black circles indicate nodes with >80% boostrap support. Orange circles indicate Acrs characterized experimentally. Scale bar indicates branch length (AU). **(D)** CRISPRi *lacZ* silencing assay using a nuclease-deficient CRISPR system, demonstrating that both AcrIB3 and AcrIB4 inhibit target DNA recognition by Cascade. **(E)** Silver stain analysis of Cas6-3xFlag (or untagged) immunoprecipitate fractions in the presence or absence of Acrs. Molecular weight marker, kDa. **(F)** Co-immunoprecipitation of His6-AcrIB3 and Cas6-3xFlag. The housekeeping sigma factor σ^A^ is shown as a non-interacting control. L, load, UB, unbound, IP, immunoprecipitate.

To investigate the mechanism by which AcrIB3 and AcrIB4 inhibit type I-B CRISPR immunity, we first tested whether they affected target DNA engagement by the Cascade complex (Fig. 3D). We designed a CRISPRi-like assay in which we deleted the nuclease *cas3* from the CRISPR locus, then targeted Cascade to a plasmid-borne *lacZ* reporter gene in *L. seeligeri* LS1. Inactivation of *cas3* ensures that target DNA bound by Cascade will not be cleaved. When we probed for *lacZ* activity by growth on plates containing X-gal, we observed CRISPR-dependent transcriptional silencing of the targeted *lacZ* gene. When we co-expressed either AcrIB3 or AcrIB4 along with Cascade, *lacZ* transcription was restored, suggesting that both anti-CRISPRs act upstream of target DNA binding by the Cascade complex, and that neither function at the level of Cas3 recruitment. Consistent with our previous observations, neither Cas5 nor Cas8b influenced target DNA recognition.

Next, we investigated whether AcrIB3 and/or AcrIB4 affect assembly of the Cascade complex. We began by constructing a type I-B CRISPR locus containing a *cas6* allele fused to a 3xFlag tag on the C-terminus. We confirmed that this fusion remained functional in interference against a plasmid with a type I-B protospacer (Fig. S13A). We then used this construct to perform anti-Flag immunoprecipitations of the Cascade complex, in the presence and absence of AcrIB3 and AcrIB4. In the absence of Acrs, the silver-stained Cas6-3xFlag immunoprecipitate fraction contained bands consistent with the molecular weights of Cas8b, Cas7, Cas6, and Cas5, none of which were present in a untagged control sample (Fig. 3E). When we co-expressed AcrIB3 or AcrIB4, each Cascade subunit remained present in the immunoprecipitate, suggesting that neither Acr impedes assembly of the type I-B Cascade complex. To test whether AcrIB3 was integrated into the complex, we fused an N-terminal his6 tag onto AcrIB3, confirmed that it was functional in inhibition of plasmid targeting by type I-B CRISPR, and performed immunoprecipitation of the Cascade complex in the presence of his6-AcrIB3 (Fig. 3F, Fig. S13b). We then analyzed the contents of the load, unbound, and immunoprecipitated fractions by immunoblotting for Cas6-3xFlag, His6-AcrIB3, and the housekeeping sigma factor σ^A^. We found that His6-AcrIB3 (but not σ^A^) strongly co-immunoprecipitated with Cas6-3xFlag, suggesting that AcrIB3 assembles into the Cascade complex. While we attempted to perform the same experiment with AcrIB4, we could not obtain a functional tagged allele.

### A Cas3 homolog inhibits type VI-A CRISPR immunity at the crRNA biogenesis stage

In addition to AcrIB3 and AcrIB4, we discovered a third Acr protein (AcrVIA2) with homology to type I-B Cas proteins (Fig. 4A). AcrVIA2 shares 24% sequence identity with the helicase-nuclease Cas3 (Fig. S12). The homology between the two proteins is centered on a shared DEAD-box helicase domain, and AcrVIA2 lacks the HD nuclease domain of Cas3. Our homology searches uncovered several true Cas3 proteins as well as 8 predicted AcrVIA2 homologs not located near a CRISPR array or *cas* gene operon, 2 of which were present on *Listeria* mobile genetic elements, while the rest were encoded in *Myoviridae* phage genomes (Fig. 4B). Again, the Acrs formed a high-confidence phylogenetic group separate from true Cas3 proteins. Surprisingly, we found that this Acr did not inhibit type I-B immunity, but instead strongly inhibited the RNA-targeting type VI-A CRISPR system (Fig. 2C). As with the two previously mentioned Cas-homolog Acrs, we confirmed that *bona fide* Cas3 possessed no inhibitory activity against Cas13 in a plasmid-targeting assay (Fig. 4C). When we mutated the AcrVIA2 DEAD box (DEFD>AAFD), we found that it lost inhibitory activity, suggesting that this domain is required for the function of AcrVIA2 (Fig. 4C). Next, we tested whether AcrVIA2 could prevent Cas13 immunity against a phage target (Fig. 4D). We infected lawns of *L. seeligeri* harboring a spacer (*spc59*) targeting the Cas13-sensitive phage ϕLS59, while co-expressing AcrVIA2 from a plasmid. While we observed a CRISPR-dependent reduction in ϕLS59 plaque formation in this system, expression of AcrVIA2 restored phage infection in the presence of Cas13 immunity. Finally, recognition of target RNA by Cas13 stimulates a non-specific *trans*-RNase activity that induces cell dormancy in *L. seeligeri*^28^. We tested whether AcrVIA2 impacts activation of Cas13 *trans* activity using a strain harboring an aTc-inducible, non-essential, non-coding RNA containing a protospacer recognized by *spc4* of the type VI-A CRISPR array (Fig. 4E). This strain is viable in the absence of target induction, but when plated on media containing aTc, exhibits a strong Cas13-dependent growth defect as a consequence of nonspecific RNase activity. In contrast, co-expression of AcrVIA2 abolished Cas13-induced dormancy, and therefore prevents cleavage of target and non-target RNA.

**Figure 4.**
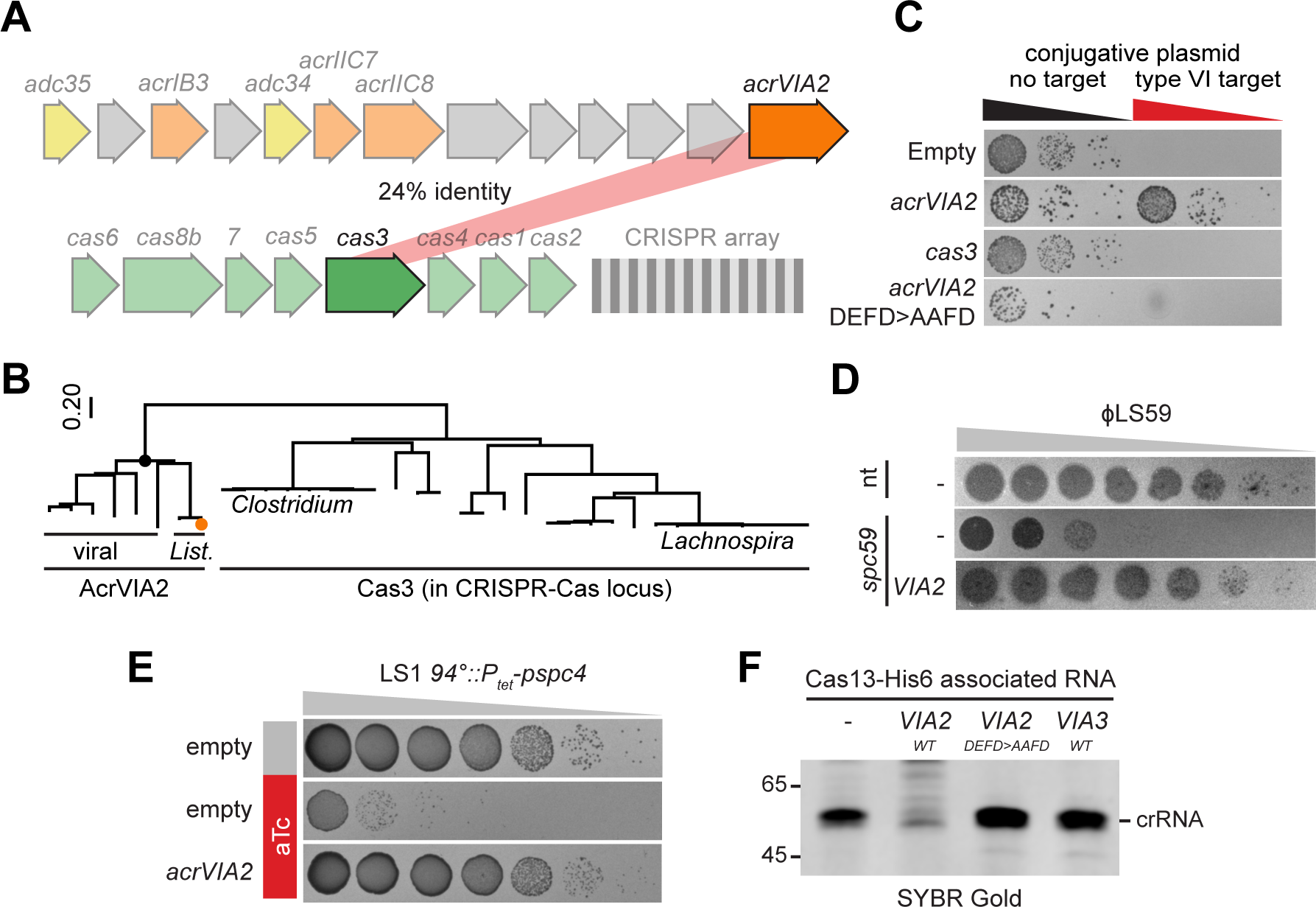
AcrVIA2 is a Cas3 homolog that inhibits type VI-A CRISPR immunity. **(A)** Schematic of genetic loci encoding AcrVIA2 (opaque orange) and type I-B CRISPR-Cas locus. Percent sequence identity between AcrVIA2 and Cas3 is noted. **(B)** Predicted phylogeny of AcrVIA2 homologs uncovered by BLAST search. Black circle indicates node with 100% bootstrap support. Orange circle indicates experimentally characterized Acr. Scale bar indicates branch length (AU). (**C**) Plasmid targeting assay demonstrating that expression of AcrVIA2, but not Cas3 or an AcrVIA2 DEAD-box mutant allele, inhibits type VI-A CRISPR immunity. **(D)** Plaque assay demonstrating that AcrVIA2 inhibits type VI-A immunity against a phage target. nt, non-targeting, spc59, spacer targeting an early lytic transcript of ɸLS59. **(E)** AcrVIA2 inhibits *trans*-RNase activity of Cas13 in vivo. Strain LS1 harboring an aTc-inducible type VI-A protospacer, plus AcrVIA2, was plated on media with or without aTc, as indicated. **(F)** The effect of AcrVIA2 on crRNA associated with affinity-purified Cas13-his6. Nucleotide molecular weight marker shown.

Next we investigated the mechanism of Cas13 inhibition by AcrVIA2. We first attempted to detect a physical interaction between both proteins. However, we were unable to detect co-immunoprecipitation of Cas13-his6 along with a partially functional AcrVIA2-3xflag allele (Fig. S14), suggesting that, unlike AcrVIA1, AcrVIA2 does not form a stable interaction with Cas13. Accordingly, we tested whether AcrVIA2 impacts the assembly of the Cas13:crRNA RNP complex. We immunoprecipitated a functionally tagged Cas13-3xFlag allele in the presence and absence of AcrVIA2, then purified RNA from the isolated protein and analyzed it by SYBR Gold staining (Fig. 4F). We detected an RNA band consistent with the mature 51 nt crRNA in the immunoprecipitated Cas13 fraction, but this band was absent from the protein purified from cells expressing AcrVIA2. Conversely, neither the AcrVIA2 DEAD-box mutant nor the unrelated protein AcrVIA3 affected Cas13-associated crRNA levels. Collectively, these results suggest that AcrVIA2 influences type VI-A crRNA processing, loading, or stability, in a mechanism that depends on its DEAD-box motif.

### Diverse viral *cas* genes reside in putative anti-defense loci

Our discovery of 3 unique Acrs homologous to type I-B Cas proteins prompted us to perform bioinformatic searches for other viral *cas* genes that might play anti-defense roles. We used 536 Cas protein query sequences to probe for *cas* genes present in the IMGVR database of high-confidence viral genomes. To enrich for putative Acrs, we then removed all hits containing nearby predicted CRISPR arrays or high-confidence *cas* gene operons. We further eliminated all genes located within 1 kb of DNA contig ends, and genes that shared greater than 90% nucleotide sequence identity with an existing hit. Ultimately, our analysis yielded 358 predicted orphan viral *cas* genes, representing components of types I, II, III, IV, and VI CRISPR-Cas systems (Fig. 5A, Table S3). The predicted hosts infected by viruses harboring orphan *cas* genes included most bacterial phyla, with *Firmicutes* and *Bacteroidota* phages being particularly abundant. We found that several of the predicted viral *cas* genes were located next to known *acr* genes or predicted anti-defense candidates from our analysis in *Listeria*, supporting the idea that some of the *cas* homologs in our dataset play anti-CRISPR roles (Fig. 5B and Fig. S15).

**Figure 5.**
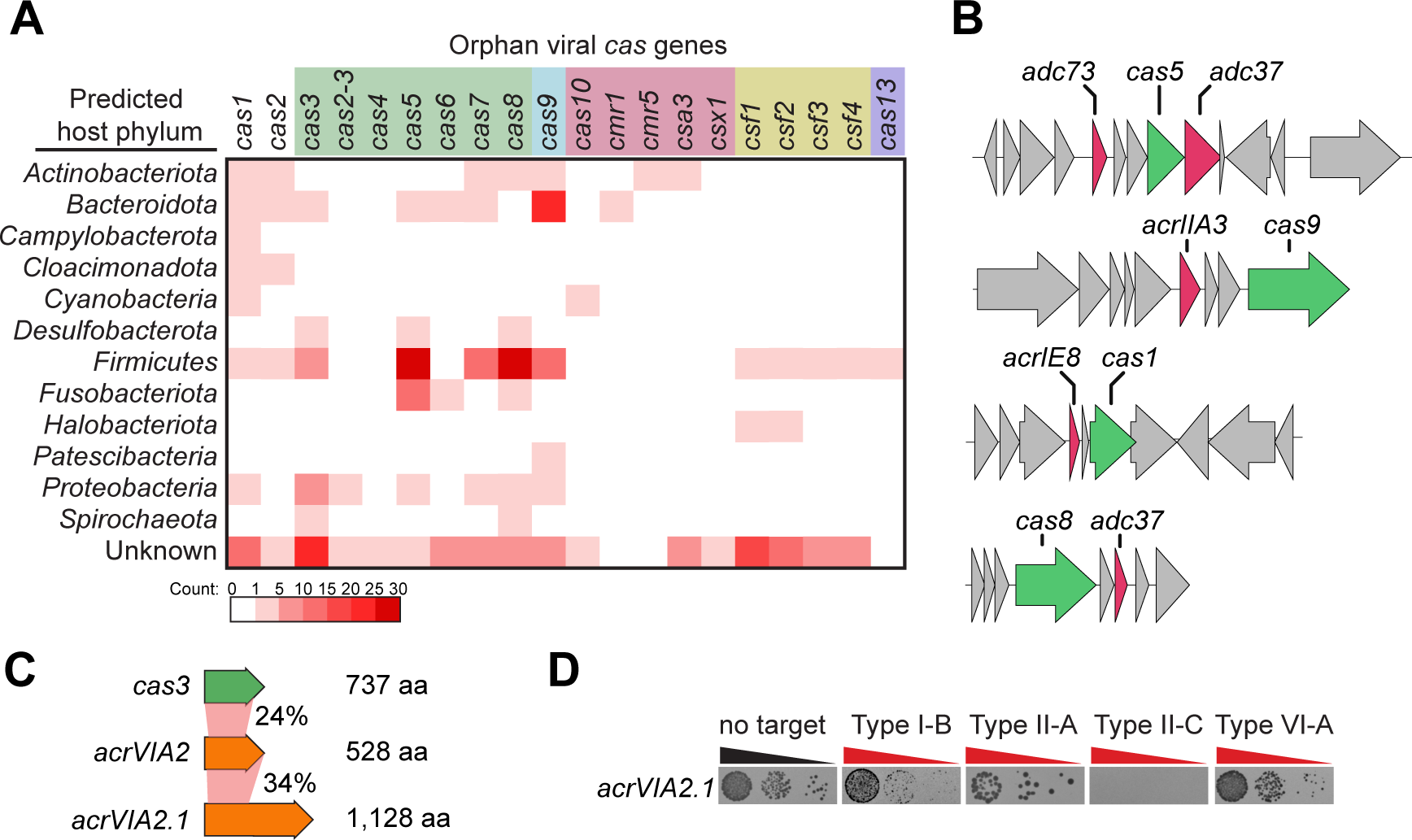
Diverse viral *cas* genes reside in putative anti-defense loci. **(A)** Frequency of orphan viral *cas* genes found in the IMGVR database, organized by Cas protein query and predicted viral host phylum. Cas queries are colored by CRISPR type (green - type I; blue - type II; pink - type III; yellow - type IV; purple - type VI). **(B)** Example loci schematics for viral *cas* genes found in the vicinity of known anti-CRISPRs or other predicted anti-defense genes. **(C)** Schematic showing percent amino acid identity between *L. seeligeri* Cas3, the indicated Acrs, and each other, along with protein lengths. (**D**) Plasmid targeting assay demonstrating the CRISPR inhibition spectrum of viral Cas3 protein.

To investigate this, we selected *acr* gene candidates with homology to *Listeria cas* genes, and tested their ability to inhibit plasmid targeting by each CRISPR type (Fig. 5C-D). Among the tested candidates was a *cas3* homolog encoded on a *Myoviridae* genome. Like AcrVIA2, this protein shared limited identity (∼24%) with the DEAD-box helicase domain of *L. seeligeri* Cas3, shared less than 40% sequence identity with AcrVIA2 (Fig. 5C), and contained no additional domains of known function. Finally, while AcrVIA2 is similar in length to Cas3, the viral Cas3 homolog identified in our bioinformatic search was over twice the size, at 1,128 amino acids. We first the viral Cas3 homolog’s ability to inhibit type VI-A CRISPR immunity against a targeted plasmid, and found that it abolished Cas13-dependent interference. Due to its homology to AcrVIA2, we refer to it as AcrVIA2.1. Next, we tested the inhibition spectrum of AcrVIA2.1 against the four *Listeria* CRISPR-Cas types (Fig. 5D). Unlike AcrVIA2, AcrVIA2.1 mediated strong inhibition of types VI-A, I-B, and II-A CRISPR interference. Thus, of all anti-CRISPRs characterized to date, AcrVIA2.1 has both the largest size and Cas protein inhibition spectrum. Collectively, our results suggest that there has been extensive *acr* evolution from *cas* genes, and that searching for orphan *cas* genes homologs in viral genomes is a useful approach to bioinformatically identify new anti-defense gene loci.

## DISCUSSION

Here we have investigated the occurrence of anti-CRISPR-mediated inhibition across a large collection of bacterial isolates, and four CRISPR-Cas types. Our results suggest the existence of widespread CRISPR antagonism present among *Listeria seeligeri* strains, which can be accounted for by 4 known and 12 previously unidentified Acr families. Three of these Acrs bear sequence identity to type I-B Cas subunits, suggesting that each Acr shares a common ancestor with its cognate Cas component. Our investigation of the mechanisms of these Acrs indicate that (i) AcrIB3 and AcrIB4 inhibit type I-B CRISPR immunity via assembly into a defective Cascade interference complex that fails to engage target DNA, and (ii) AcrVIA2 inhibits type VI-A CRISPR immunity by blocking the processing, loading, or stability of Cas13-associated crRNA. To investigate the generality of Acr evolution from Cas proteins, we probed the IMGVR database for the existence of orphan viral *cas* genes. We uncovered hundreds of examples of viral *cas* genes that were not associated with a CRISPR array or complete *cas* gene operon, instead residing near putative anti-defense genes. We experimentally confirmed that at least one of these genes (AcrVIA2.1) exhibits exceptionally broad-spectrum inhibition of CRISPR-Cas immunity in *L. seeligeri*. In addition to uncovering numerous anti-CRISPR proteins that could potentiate phage therapy or gene editing safety, our findings demonstrate that diverse viruses have co-opted *cas* genes for CRISPR antagonism, and provide a new strategy for the unbiased identification of counter-defense genes in prokaryotes.

Our results raise several questions regarding the evolutionary trajectories that could convert a host-encoded *cas* gene to a phage-encoded *acr.* First, how do phages capture *cas* genes? One possibility is via imprecise excision of temperate phages integrated near CRISPR-Cas loci. During induction of such prophages, *cas* genes could occasionally be packaged into viral capsids along with the phage genome. Varble and colleagues^30^ recently demonstrated that some *Streptococcus* phages integrate directly into the degenerate repeats of type II-A CRISPR arrays, and can sometimes capture and mobilize spacer sequences. It remains to be seen whether such a mechanism could also promote viral capture of whole *cas* genes or fragments thereof. Once a *cas* gene is integrated into a phage genome, it may not immediately play a role in CRISPR antagonism. Instead, viral *cas* genes might stimulate CRISPR immunity to play a protective role for lysogenized hosts that could otherwise be infected by a second phage. Next, how is a viral *cas* gene exapted into an anti-CRISPR? Because Cas proteins naturally make interactions with other Cas proteins, crRNA, and target nucleic acids, they are well-poised to evolve into inhibitors that block CRISPR immunity. Any phage-encoded Cas protein that interacts with two or more components of the CRISPR RNP might develop inhibitory activity by simply losing one of these interactions while maintaining another, resulting in a faulty Cas subunit that inactivates immunity. One benefit of this strategy (as compared to non-Cas anti-CRISPRs) is that it may be difficult for CRISPR systems to evolve resistance against such inhibitors, since they resemble the very Cas components used for immunity. Finally, our results raise the possibility that subunits of anti-phage immune systems beyond CRISPR may also serve as raw material for counter-defense evolution.

In this study, we uncovered a total of 12 anti-CRISPR families present in *Listeria* prophages and mobile genetic elements. Residing beside these *acr* genes were 64 additional anti-defense candidate genes, 26 of which exhibited no detectable CRISPR inhibition in our assay (Table S2). While some of these genes may serve other functions, their frequent co-occurrence with and proximity to *acr* genes suggests that many could play an anti-defense role, possibly against one or more of the other anti-viral defense systems found in *Listeria spp.* Indeed, recent studies have uncovered examples of viral inhibitors of CBASS, Pycsar, Thoeris, Gabija, and Hachiman defenses^31–34^.

Though CRISPR-Cas systems are abundant in *Listeria spp.*, our functional screen revealed that most are inhibited by endogenous Acrs. Such frequent inhibition likely provides a selective pressure to acquire new diverse immune systems not susceptible to existing Acrs. For example, while we observed inhibition of the highly abundant type I-B CRISPR in 77% of the tested *L. seeligeri* strains, the less common type VI-A system was only inhibited in 29% of strains. If inhibition is widespread, why are CRISPR systems retained by the host? On the contrary, recent evidence suggests that prophage-encoded Acrs promote retention of host CRISPR-Cas systems, by preventing autoimmune cleavage of targets within the integrated prophage^35^. Maintenance of functional CRISPR immunity despite the presence of Acrs could provide a fitness benefit in the event that the host becomes cured of the prophage or MGE harboring *acr* genes. In total, our findings represent an example of the diversity of evolved interactions in the ongoing phage-host arms race.

## Supporting information

Table S1 Strains Plasmids and Oligos used in this study

Table S2 Experimentally verified Acrs

Table S3 Predicted orphan viral cas genes

## ACKNOWLEDGEMENTS

We are grateful to all members of the Meeske lab for advice and encouragement, members of the Guo and Mitchell labs for helpful discussions. This work in the AJM lab was supported by the NIH [R35GM142460, S10OD026741], NSF [FAIN2235762] and the University of Washington Royalty Research Foundation. SRM is supported by the Jane Coffin Childs Memorial Fund for Medical Research. The work in the JBD lab was supported by the National Institutes of Health (R01GM127489) and funding from the Defense Advanced Research Projects Agency (DARPA) award HR0011-17-2-0043. The views, opinions and/or findings expressed are those of the authors and should not be interpreted as representing the official views or policies of the Department of Defense or the US Government, and the DARPA Safe Genes program (HR0011-17-2-0043).

## AUTHOR CONTRIBUTIONS

The project was conceived by AJM. Experiments were performed by MAK, EMS, AK, MCW, SRM, and AJM. Bioinformatic analysis was conducted by MAK, MJ, JB-D, and AJM. MJ was supervised by JB-D. The manuscript was written by MAK and AJM, all authors contributed to manuscript editing.

## METHODS

### Bacterial strains and growth conditions

*L. seeligeri* strains were cultured in Brain Heart Infusion (BHI) medium at 30°C. Where appropriate, BHI was supplemented with the following antibiotics for selection: nalidixic acid (50 μg/mL), chloramphenicol (10 μg/mL), erythromycin (1 μg/mL), or kanamycin (50 μg/mL). For cloning, plasmid preparation and conjugative plasmid transfer, *E. coli* strains were cultured in Lysogeny Broth (LB) medium at 37 °C. Where appropriate, LB was supplemented with the following antibiotics: ampicillin (100 μg/mL), chloramphenicol (25 μg/mL), kanamycin (50 μg/mL). For conjugative transfer of *E. coli* – *Listeria* shuttle vectors, plasmids were purified from Turbo Competent *E. coli* (New England Biolabs) and transformed into the *E. coli* conjugative donor strains SM10 λ*pir* or S17 λ*pir.* For a list of strains used in this study, see Table S1.

### Plasmid construction and preparation

All genetic constructs for expression in *L. seeligeri* were cloned into the following three compatible shuttle vectors, each of which contains an origin of transfer sequence for mobilization by transfer genes of the IncP-type plasmid RP4. These transfer genes are integrated into the genome of the *E. coli* conjugative donor strains SM10 λ*pir* or S17 λ*pir.* All plasmids used in this study, along with details of their construction can be found in Table S1.

pPL2e: ectopically integrating plasmid conferring chloramphenicol resistance in *E. coli* and erythromycin resistance in *Listeria*; integrates into the tRNA^Arg^ locus in the *L. seeligeri* chromosome^29^.

pAM8: *E. coli-Listeria* shuttle vector conferring ampicillin resistance in *E. coli* and chloramphenicol resistance in *Listeria*^27^.

pAM326: *E. coli-Listeria* shuttle vector conferring kanamycin resistance in *E. coli* and *Listeria*^12^.

Mobilizable CRISPR-Cas systems were constructed by cloning the type I-B, II-A, II-C, and VI-A CRISPR-Cas loci into pPL2e, each equipped with a spacer matching a target plasmid. Target plasmids were derived from pAM8. In the case of type II-A, one variant of the CRISPR-Cas plasmid harbored a spacer targeting a protospacer region on pAM8 followed by an NGG PAM, and a separate CRISPR plasmid harbored a non-targeting spacer. The same approach was taken for type II-C, except the protospacer region was followed by an NNGCAA PAM. For types I-B and VI-A, naturally occurring spacers were used in the CRISPR plasmid, and matching protospacers were inserted into pAM8. The type I-B protospacer was preceded by a 5’ CCN PAM. The type VI-A protospacer was inserted into a transcribed region in the 3’ UTR of the chloramphenicol resistance gene of pAM8.

Putative anti-CRISPR constructs were assembled by cloning into NcoI/EagI digested pAM551, which is derived from pAM326 and contains an aTc-inducible P_tet_ promoter.

### E. coli – L. seeligeri conjugation

All genetic constructs for expression in *L. seeligeri* were introduced by conjugation with *E. coli* donor strains SM10 λ*pir* or S17 λ*pir.* 100 μL of each donor and recipient culture were diluted into 10 mL of BHI medium and concentrated on a 0.45 µm porosity filter disk using vacuum filtration. Filter disks were laid onto BHI agar supplemented with oxacillin (8 μg/mL for pPL2e or pAM326 derived plasmids and 128 μg/mL for pAM8 derived plasmids) which weakens the cell wall and enhances conjugation, then incubated at 37°C for 4 hours. Cells were resuspended in 2 mL BHI, serially diluted, and transconjugants were selected on BHI medium containing 50 μg/mL nalidixic acid (which kills donor *E. coli* but not recipient *L. seeligeri)* in addition to the appropriate antibiotic for plasmid selection. Transconjugants were isolated after 2-3 days of incubation at 30°C.

### Phylogenetic tree construction

To reconstruct Acr phylogeny, query Acr proteins were searched against the BLAST nr database^36^ using an E-value cutoff of 5×10^-3^ (for AcrIB4) or 1×10^-4^ (for all other Acrs). The top 250 hits were aligned using T-Coffee^37^. For AcrIB4, only the C-terminal 90 amino acids were included for alignment, as this is the region with shared homology between AcrIB4 and the much larger Cas8b. Phylogenetic trees and bootstrap values were calculated using MEGA (v11)^38^, using the neighbor-joining method with 1000 bootstrap replications.

### Co-immunoprecipitation

*L. seeligeri* harboring FLAG-tagged and/or His6-tagged proteins was cultured in 30 mL BHI to saturation, then pelleted by centrifugation. Cells were resuspended in 1.5 mL lysis buffer containing 50 mM HEPES pH 7, 150 mM NaCl, 5 mM MgCl_2_, 5% glycerol, 1 mM PMSF, and 2 mg/mL lysozyme, then incubated at 37°C for 20 min. Lysis was performed by sonication, then insoluble material was pelleted by centrifugation at 15,000 rpm for 10 min. The clarified supernatants were sampled (load fraction), then applied to 40 µL of buffer-equilibrated M2 anti-Flag antibody affinity resin (Sigma-Aldrich) and incubated at 4°C for 2 h. Flag resin was pelleted by centrifugation at 1,000 rpm for 1 min, and the supernatant was sampled (unbound fraction). Flag resin was washed three times for 5 min each with 1 mL wash buffer (50 mM HEPES pH 7, 150 mM NaCl, 5 mM MgCl_2_, 5% glycerol). Finally, the immunoprecipitated fraction was eluted with 40 µL of 0.1 mg/mL 3xFlag peptide (Sigma-Aldrich) at room temperature. All samples were denatured by dilution in 2x Laemmli sample buffer containing 4% SDS and 10% beta-mercaptoethanol. Load, unbound, and IP fractions were analyzed by immunoblot using anti-Flag (Sigma-Aldrich), anti-His6 (Genscript), and anti-σ^A^ (gift of David Rudner, Harvard Medical School) antibodies. Silver staining was performed on 12 µL of each immunoprecipitate sample, using the Pierce Silver Staining Kit (Thermo Fisher) according to the manufacturer’s instructions.

### Analysis of Cas13-associated crRNA

*L. seeligeri* cultures harboring *cas13-his6* and/or *acrVIA2* were grown to saturation. 50 mL culture was harvested, pelleted at 4300 rpm and frozen at −80°C. Pellets were resuspended in ice-cold lysis buffer (50 mM HEPES pH7.0, 150 mM NaCl, 5mM MgCl2, 10 mM imidazole, 1 mg/mL lysozyme, 1 mM phenylmethylsulfonylfluoride, 5% glycerol) and lysed by sonication. Lysate was centrifuged at 15,000 rpm for 15 minutes at 4°C. Soluble material was batch bound for 2 hours with 50uL of Ni-NTA HisBind Resin. Resin was then washed three times with 1 mL wash buffer (50 mM HEPES pH7.0, 150 mM NaCl, 5mM MgCl2, 10 mM imidazole 5% glycerol) and eluted with wash buffer supplemented with 250 mM imidazole. RNA was purified using the Direct-zol RNA miniprep kit (Zymo Research). Samples were resolved by denaturing 15% TBE-Urea PAGE, stained with SYBR Gold according to the manufacturer’s instructions, and imaged on an Azure Biosystems Azure 600 imager.

### Phage propagation

All phage infections were performed in BHI medium supplemented with 5 mM CaCl2. To generate phage lysates, existing phage stocks were diluted to single plaques on a lawn of *L. seeligeri* LS1 *ΔRM1 ΔRM2* and a single plaque was purified twice to ensure homogeneity. 5 mL of cell culture was infected with phage at MOI 0.1, OD 0.1 and the infection proceeded overnight. The lysate was centrifuged for 20 minutes at 4,000 rpm and the supernatant was filtered using a 0.45 μm pore syringe filter.

### Bioinformatic identification of viral *cas* genes

The IMGVR7.1 database of high-confidence viral genomes^39^ was probed for sequences with homology to 536 Cas protein query sequences, representing all known CRISPR subtypes^13^. Each query was searched against IMGVR7.1 using tblastn^36^ with an E-value cutoff of 1×10^-4^. 20 kb of genomic sequence flanking each hit gene was retrieved using bedtools^40^, and hits were deduplicated using genometools sequniq^41^. Hit genomic regions were analyzed for *bona fide* CRISPR-Cas systems using CRISPRCasTyper^42^, and all hits containing either predicted CRISPR arrays or *cas* gene operons were removed from analysis. Hits were further filtered to remove any *cas* genes located within 1 kb of a contig end, and hits sharing greater than 90% nucleotide sequence identity were collapsed using T-Coffee seq_reformat^37^. Finally, the IMGVR7.1 database was probed as above for homologs of known Acrs, anti-restriction-modification^43^, anti-Hachiman^33^, anti-Gabija, and anti-Thoeris genes, and hits within 10 kb of a predicted *cas* gene were tabulated. The UViG identifier for each hit was used to retrieve predicted host phylogeny from IMGVR. For gene loci diagrams, ORFs were predicted with prokka^44^ and diagrams were generated with Clinker^45^.

---

**Figure S1.**
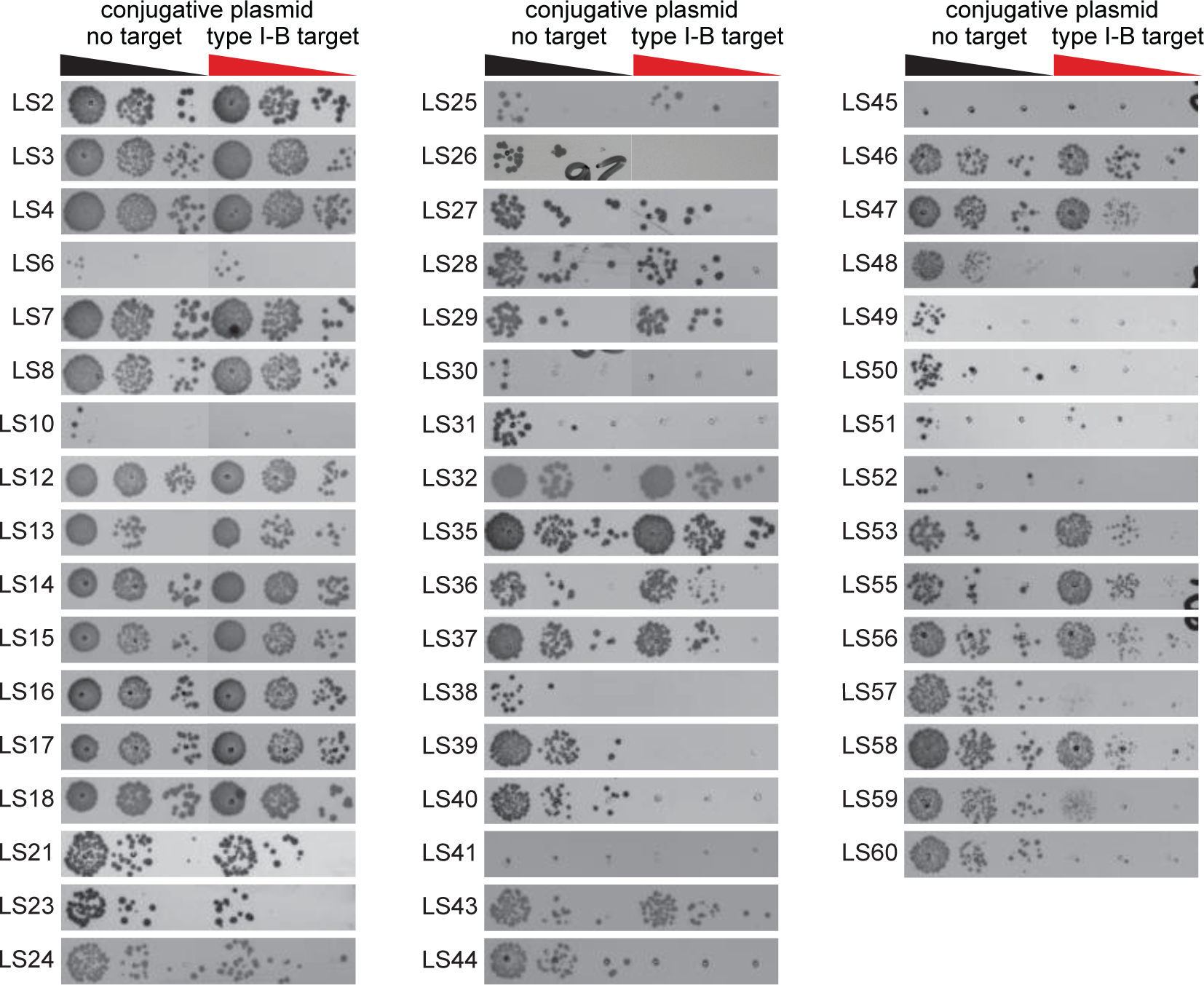
Variation in *L. seeligeri* strain background affects type I-B CRISPR-Cas immunity. Plasmid targeting assay in which the indicated L. seeligeri strains were first transformed with a chromosomally integrated type I-B CRISPR-Cas system equipped with a spacer targeting a conjugative plasmid, then challenged with either a non-target plasmid (left columns) or plasmid containing a target protospacer (right columns).

**Figure S2.**
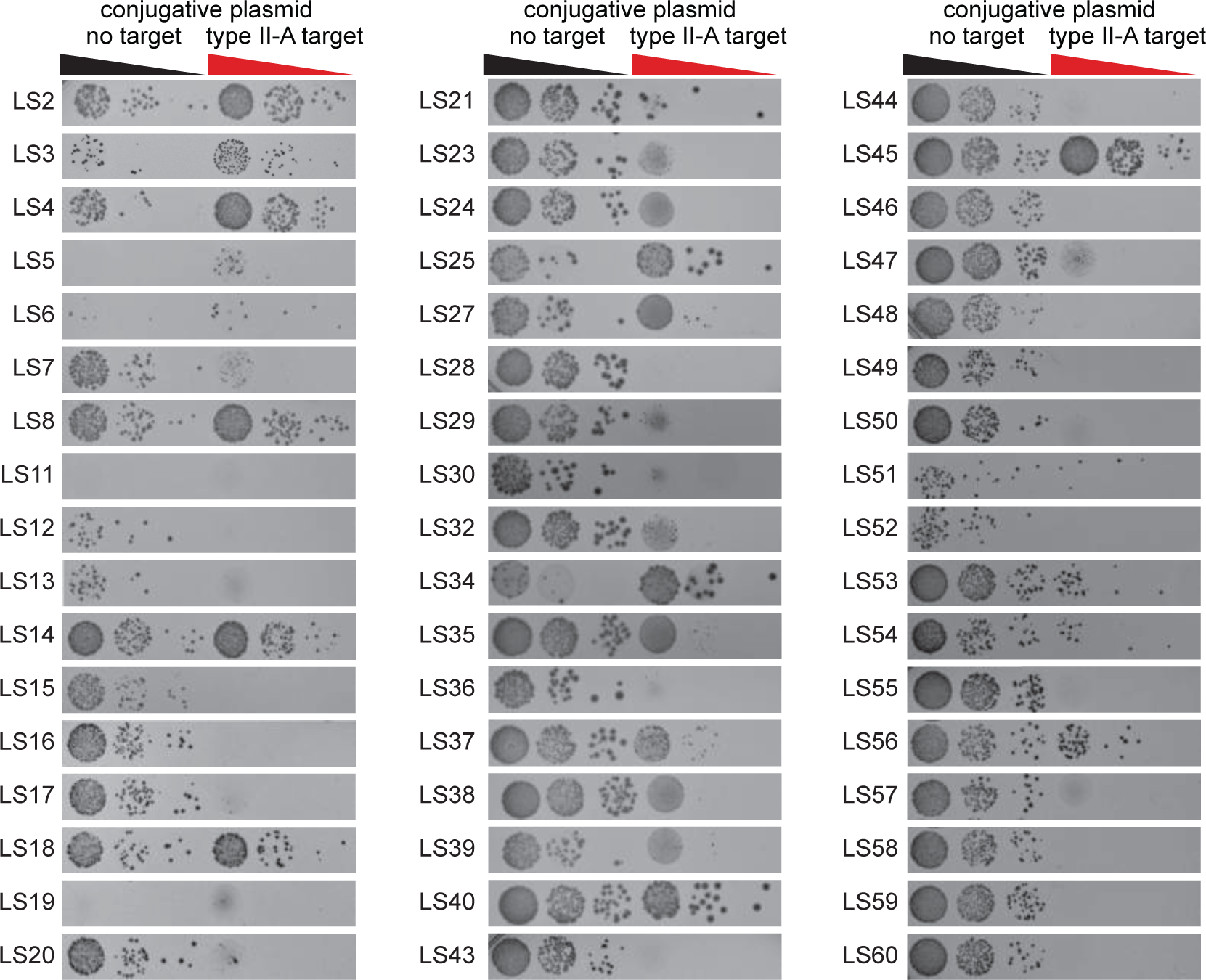
Variation in *L. seeligeri* strain background affects type II-A CRISPR-Cas immunity. Plasmid targeting assay in which the indicated L. seeligeri strains were first transformed with a chromosomally integrated type II-A CRISPR-Cas system equipped with a spacer targeting a conjugative plasmid, then challenged with either a non-target plasmid (left columns) or plasmid containing a target protospacer (right columns).

**Figure S3.**
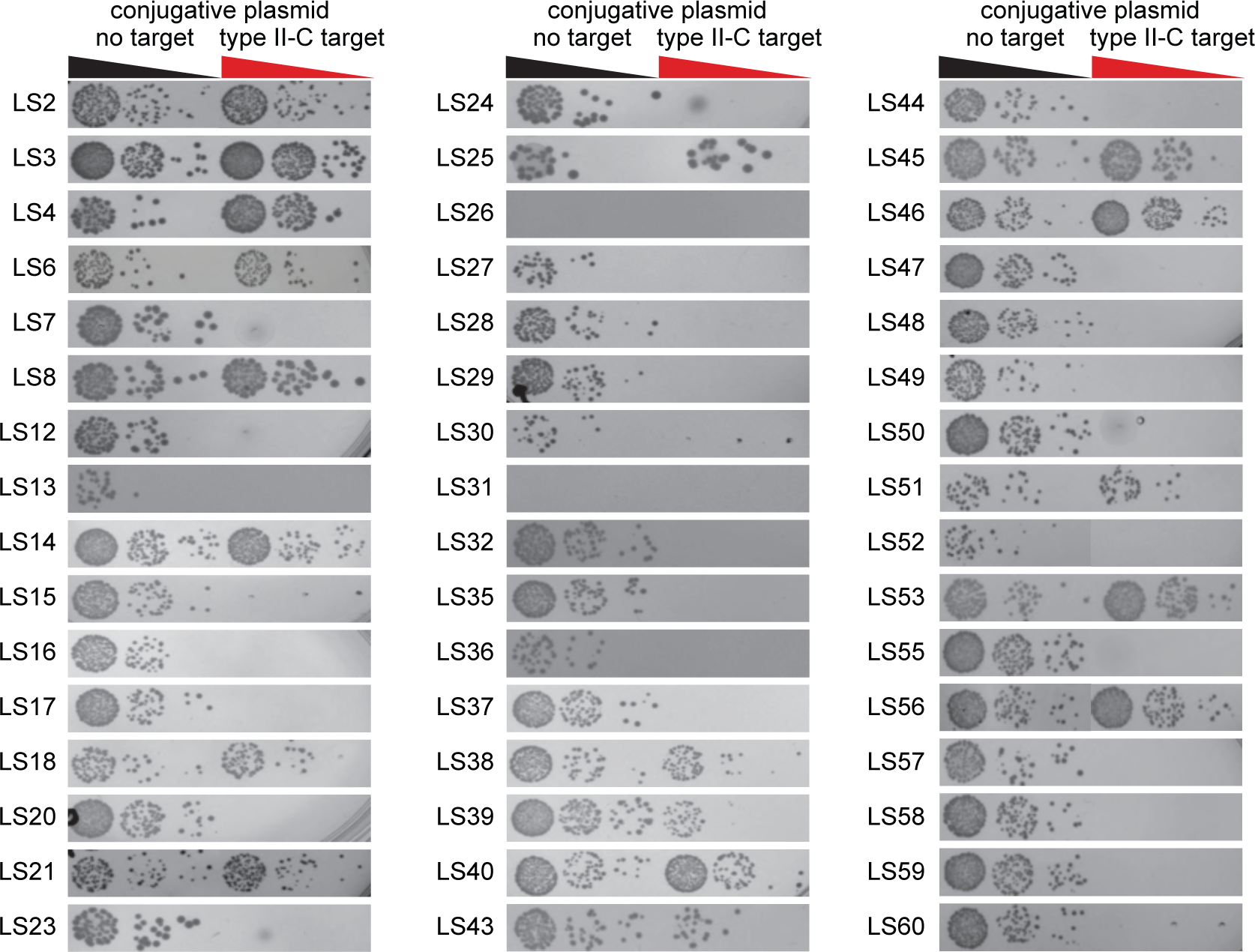
Variation in *L. seeligeri* strain background affects type II-C CRISPR-Cas immunity. Plasmid targeting assay in which the indicated L. seeligeri strains were first transformed with a chromosomally integrated type II-C CRISPR-Cas system equipped with a spacer targeting a conjugative plasmid, then challenged with either a non-target plasmid (left columns) or plasmid containing a target protospacer (right columns).

**Figure S4.**
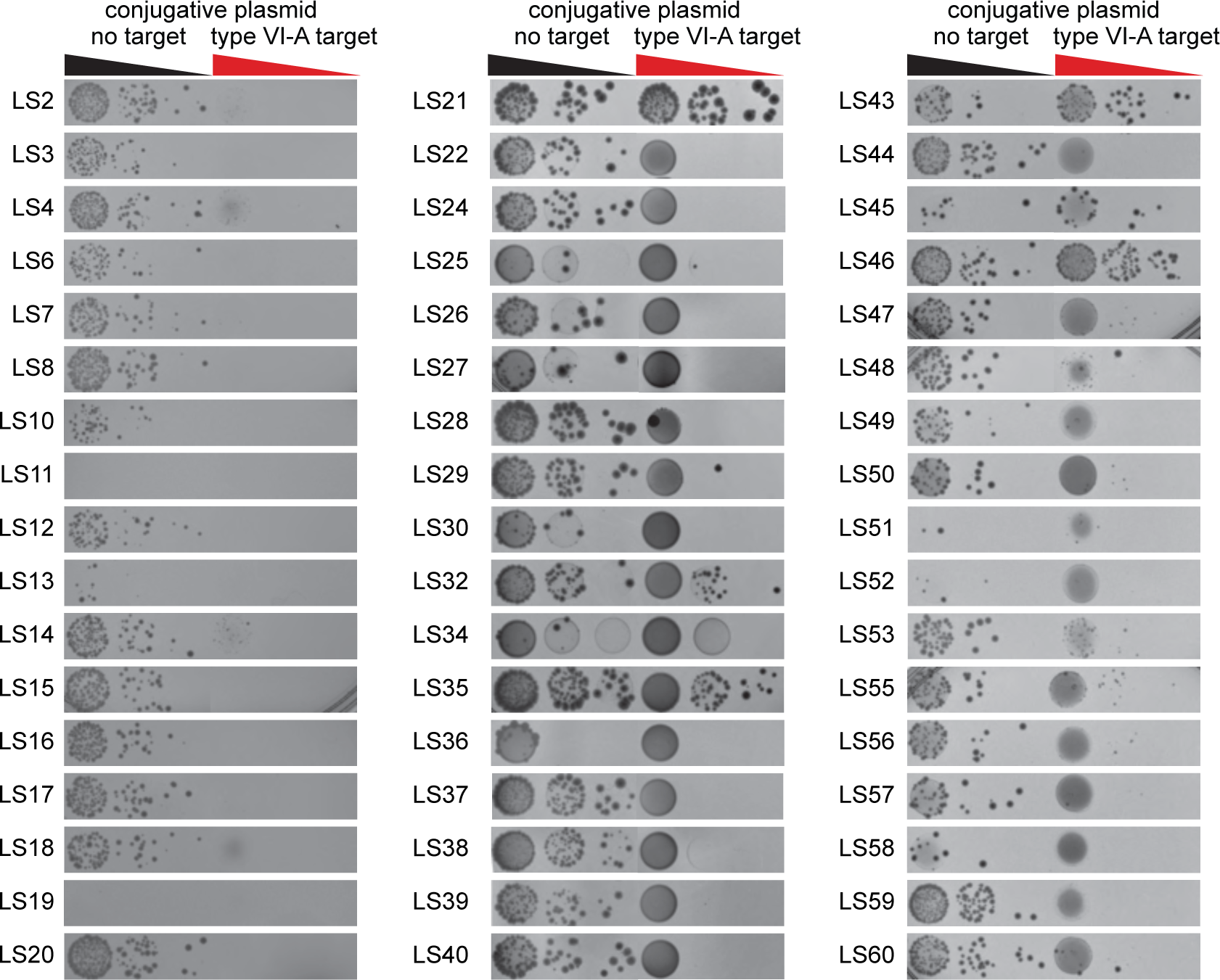
Variation in *L. seeligeri* strain background affects type VI-A CRISPR-Cas immunity. Plasmid targeting assay in which the indicated L. seeligeri strains were first transformed with a chromosomally integrated type VI-A CRISPR-Cas system equipped with a spacer targeting a conjugative plasmid, then challenged with either a non-target plasmid (left columns) or plasmid containing a target protospacer (right columns).

**Figure S5.**
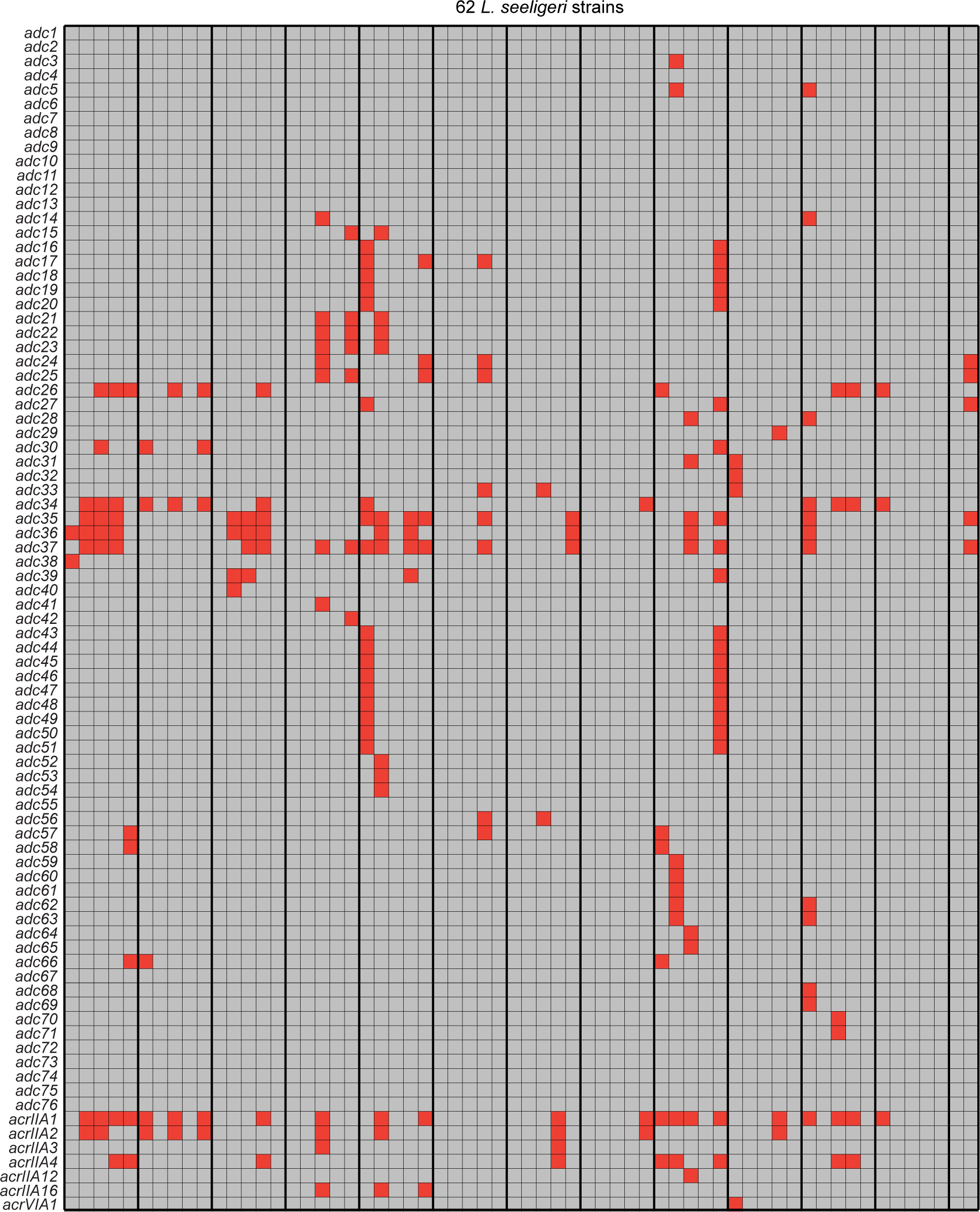
Anti-defense candidate (*adc*) gene occurrence across 62 strains of *L. seeligeri*. Each row correponds to either a known anti-CRISPR gene or a particular anti-defense candidate gene identified as frequently encoded nearby *acr* genes or nearby other well-established anti-defense canddiates. Each column corresponds to an individual *L. seeligeri* strain genome. Filled red boxes indicate occurence of a putative anti-defense gene in a particular strain.

**Figure S6.**
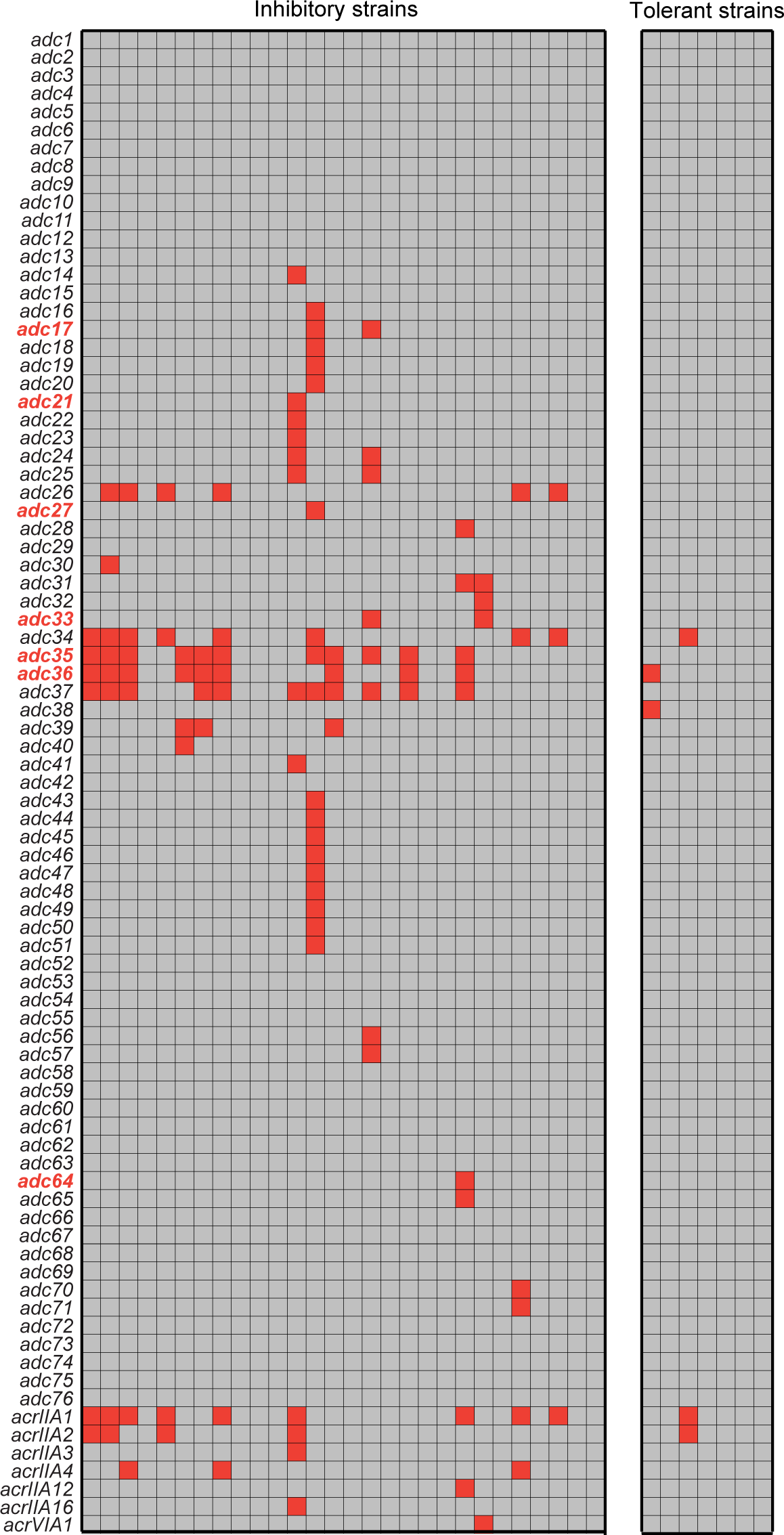
Anti-defense candidate (*adc*) gene occurrence among *L. seeligeri* strains that inhibit (or tolerate) type I-B CRISPR immunity. Each row correponds to either a known anti-CRISPR gene or a particular anti-defense candidate gene identified as frequently encoded nearby *acr* genes or nearby other well-established anti-defense canddiates. Each column corresponds to an individual *L. seeligeri* strain genome. The group of columns on the left indicate strains that inhibited type I-B CRISPR immunity in the plasmid targeting assay shown in Fig. 1, while the group on the right tolerated type I-B immunity. Filled red boxes indicate occurence of a putative anti-defense gene in a particular strain. Gene names in red indicate experimentally validated type I-B Acrs from this study.

**Figure S7.**
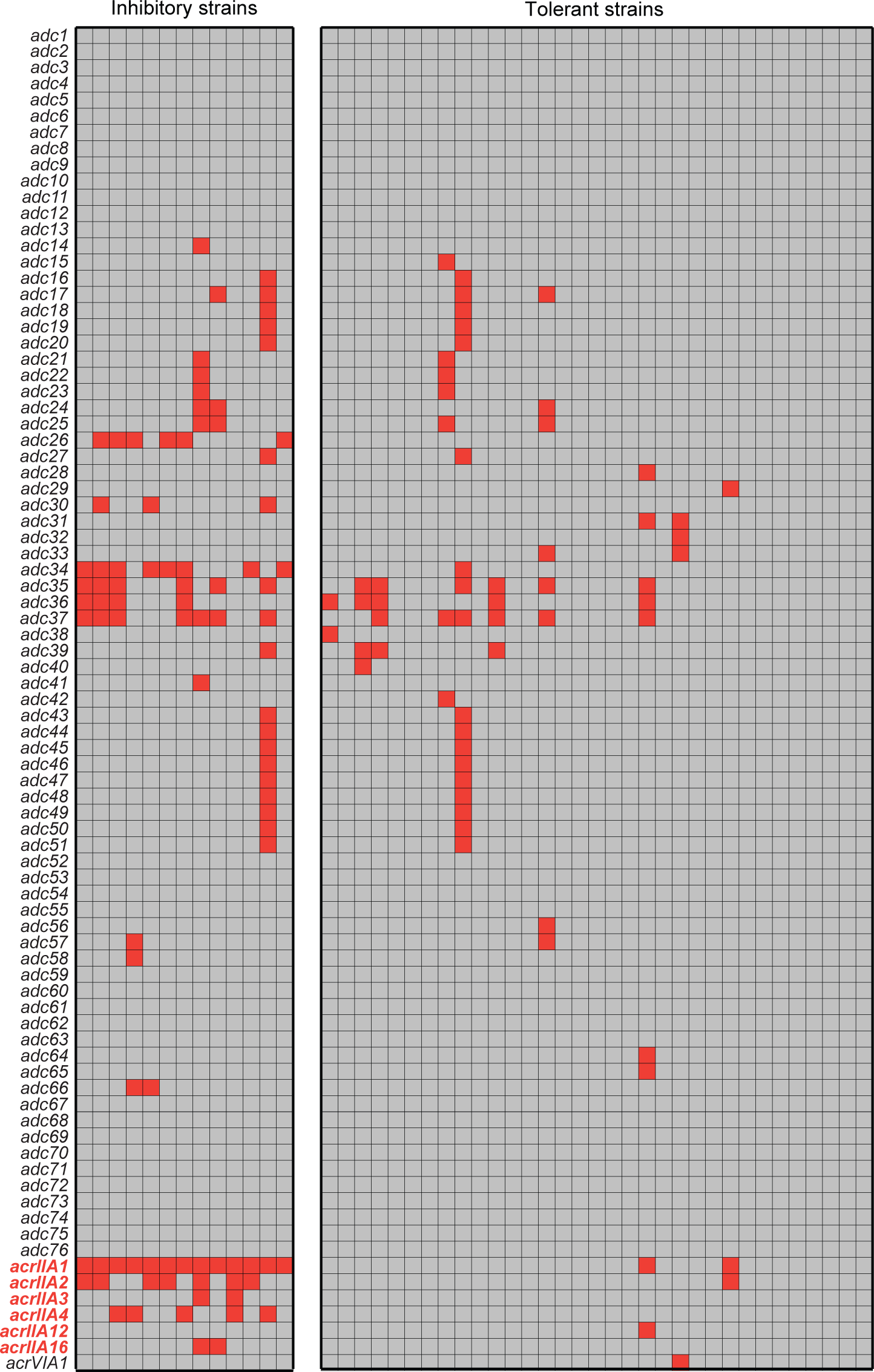
Anti-defense candidate (*adc*) gene occurrence among *L. seeligeri* strains that inhibit (or tolerate) type II-A CRISPR immunity. Each row correponds to either a known anti-CRISPR gene or a particular anti-defense candidate gene identified as frequently encoded nearby *acr* genes or nearby other well-established anti-defense canddiates. Each column corresponds to an individual *L. seeligeri* strain genome. The group of columns on the left indicate strains that inhibited type II-A CRISPR immunity in the plasmid targeting assay shown in Fig. 1, while the group on the right tolerated type II-A immunity. Filled red boxes indicate occurence of a putative anti-defense gene in a particular strain. Gene names in red indicate experimentally validated type II-A Acrs.

**Figure S8.**
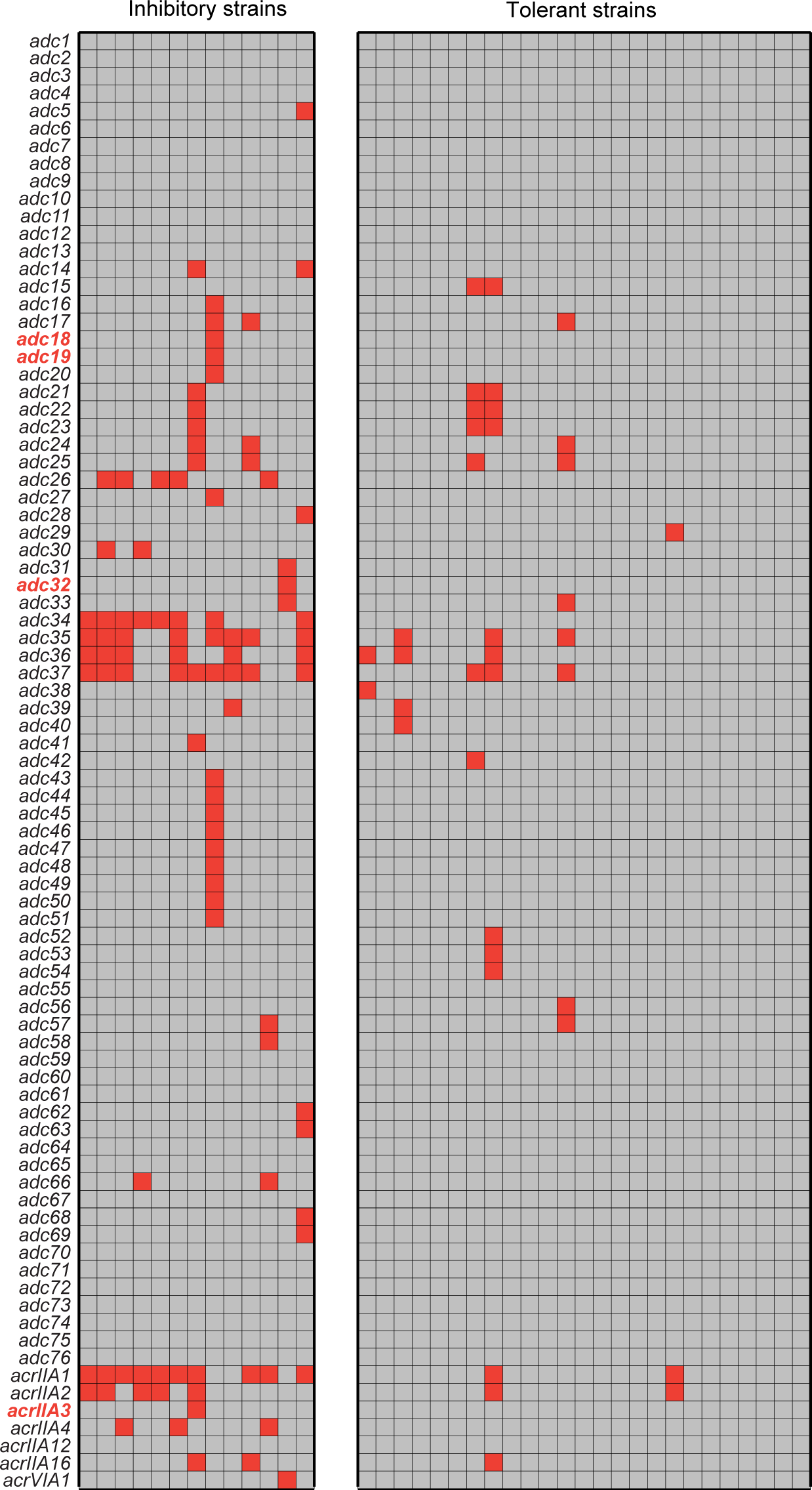
Anti-defense candidate (*adc*) gene occurrence among *L. seeligeri* strains that inhibit (or tolerate) type II-C CRISPR immunity. Each row correponds to either a known anti-CRISPR gene or a particular anti-defense candidate gene identified as frequently encoded nearby *acr* genes or nearby other well-established anti-defense canddiates. Each column corresponds to an individual *L. seeligeri* strain genome. The group of columns on the left indicate strains that inhibited type II-C CRISPR immunity in the plasmid targeting assay shown in Fig. 1, while the group on the right tolerated type II-C immunity. Filled red boxes indicate occurence of a putative anti-defense gene in a particular strain. Gene names in red indicate experimentally validated type II-C Acrs from this study.

**Figure S9.**
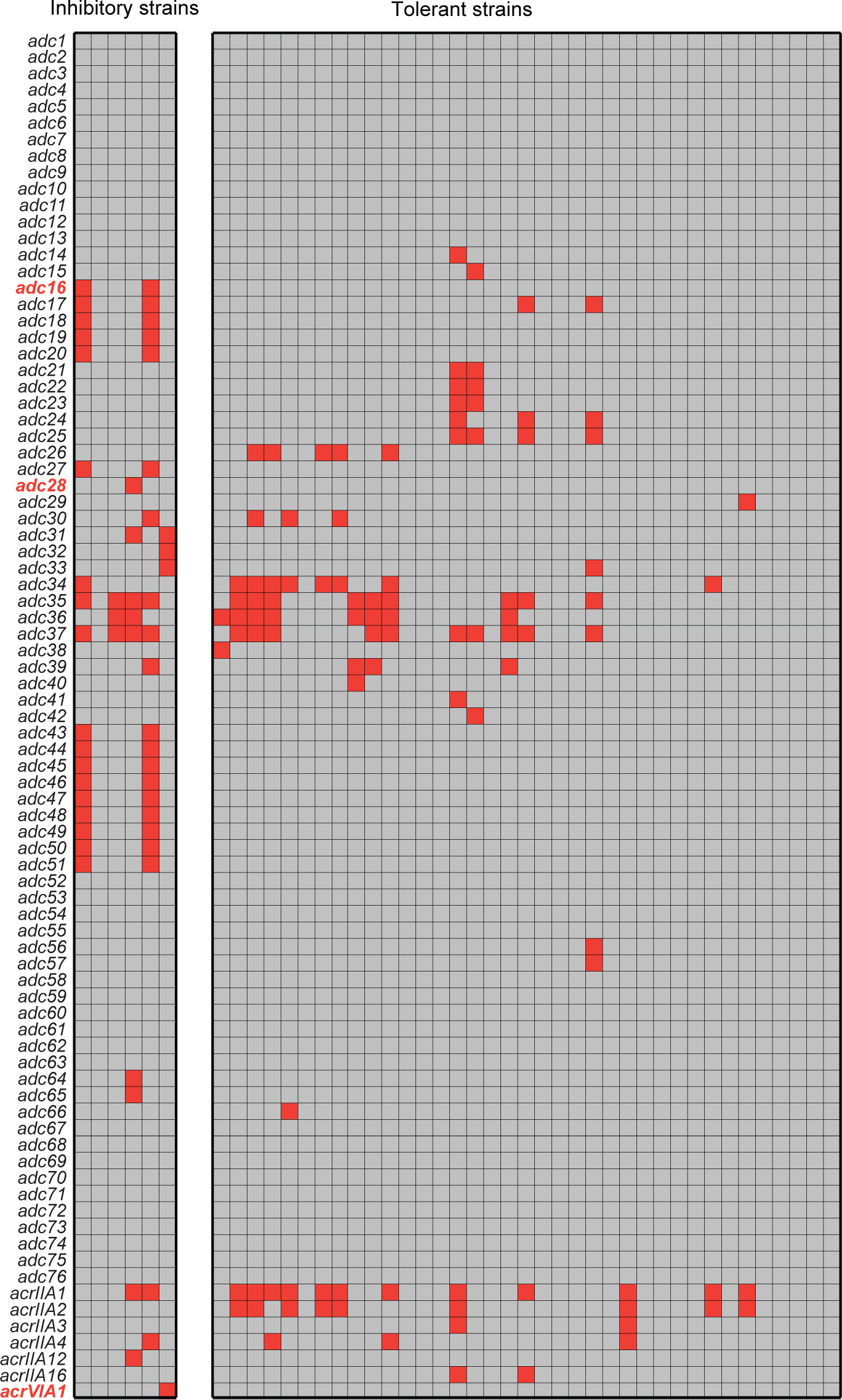
Anti-defense candidate (*adc*) gene occurrence among *L. seeligeri* strains that inhibit (or tolerate) type VI-A CRISPR immunity. Each row correponds to either a known anti-CRISPR gene or a particular anti-defense candidate gene identified as frequently encoded nearby *acr* genes or nearby other well-established anti-defense canddiates. Each column corresponds to an individual *L. seeligeri* strain genome. The group of columns on the left indicate strains that inhibited type VI-A CRISPR immunity in the plasmid targeting assay shown in Fig. 1, while the group on the right tolerated type VI-A immunity. Filled red boxes indicate occurence of a putative anti-defense gene in a particular strain. Gene names in red indicate experimentally validated type VI-A Acrs from this study.

**Figure S10.**
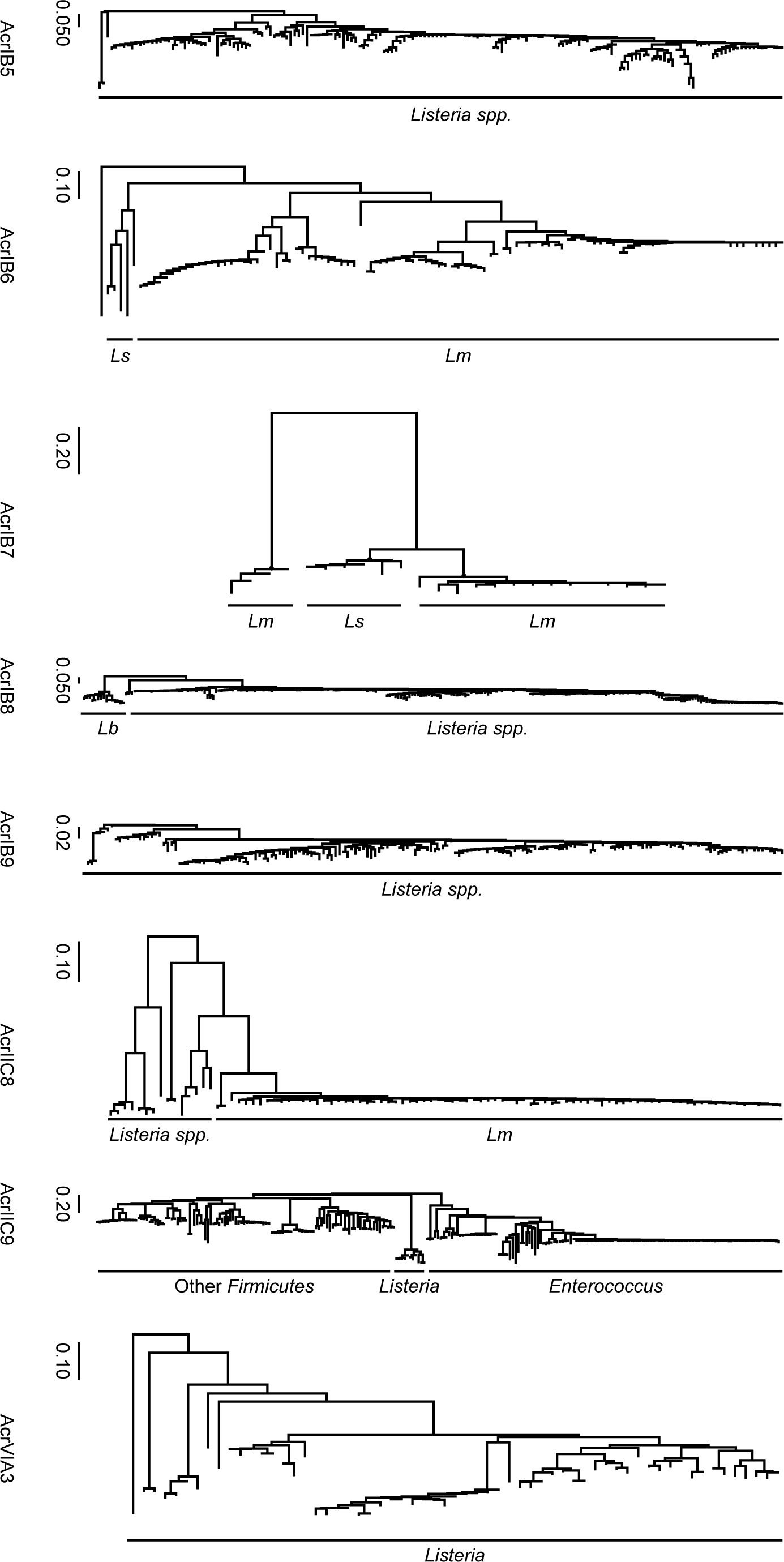
Predicted phylogeny of newly discovered type I-B, II-C, and VI-A anti-CRISPR proteins. Predicted phylogeny of Acr homologs uncovered by BLAST search. Scale bar indicates branch length (AU). See Fig. 3 for predicted phylogeny of AcrIB3 and AcrIB4 homologs. See Fig. 4 for predicted phylogeny of AcrVIA2 homologs.

**Figure S11.**
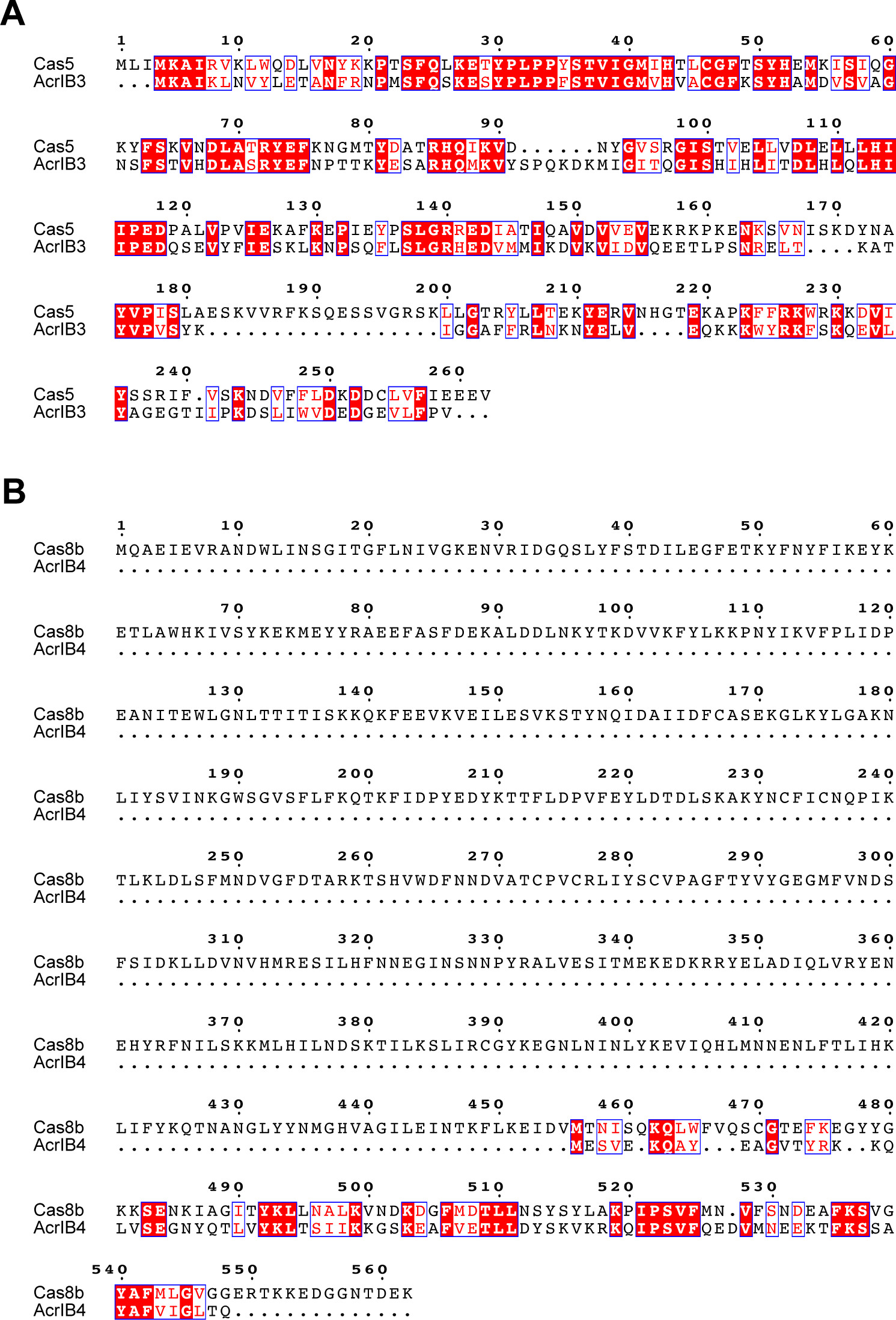
Sequence alignment of type I-B Acrs and type I-B Cas proteins. **(A)** Sequence alignment of AcrIB3 and Cas5 from *L. seeligeri* strain LS1. Identical residues are highlighted in red, while similar residues are in red text with blue outlines. **(B)** As for (A), but for AcrIB4 and Cas8b.

**Figure S12.**
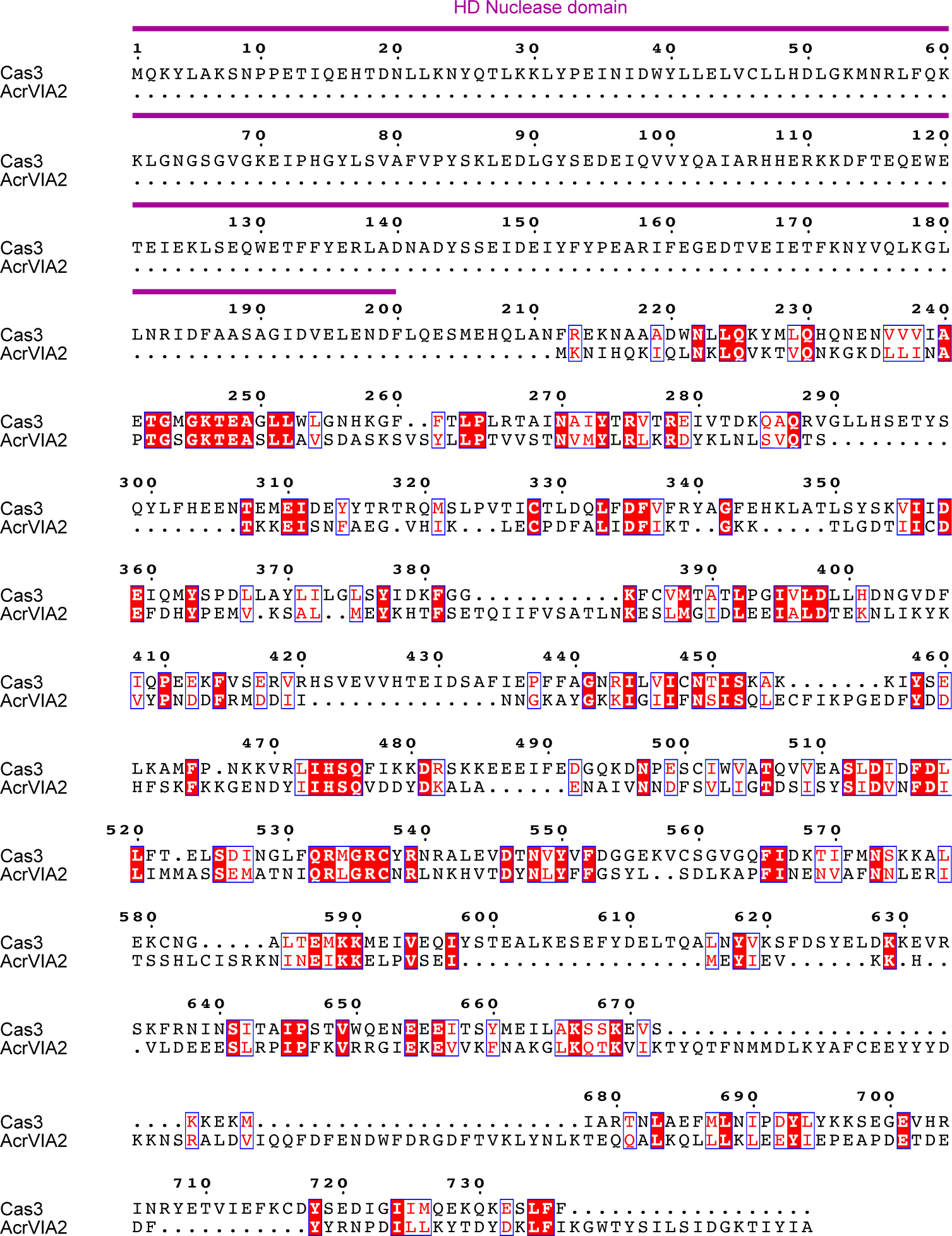
Sequence alignment of AcrVIA2 and the type I-B helicase-nuclease Cas3. Sequence alignment of AcrVIA2 and Cas3 from *L. seeligeri* strain LS1. Identical residues are highlighted in red, while similar residues are in red text with blue outlines.

**Figure S13.**
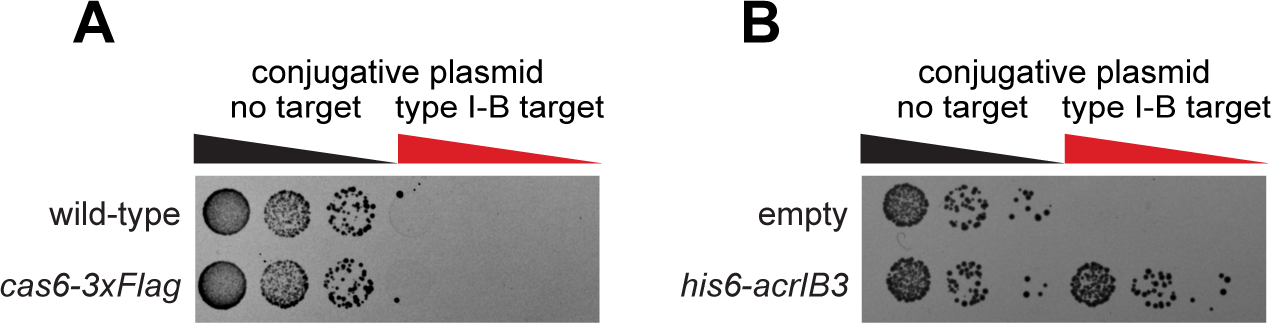
Functionality of affinity-tagged Cas proteins. **(A)** Cas6-3xFlag functions in immunity against a plasmid containing a protospacer recognized by the type I-B CRISPR system. **(B)** His6-AcrIB3 functions in inhibition of type I-B CRISPR immunity in the plasmid targeting assay.

**Figure S14.**
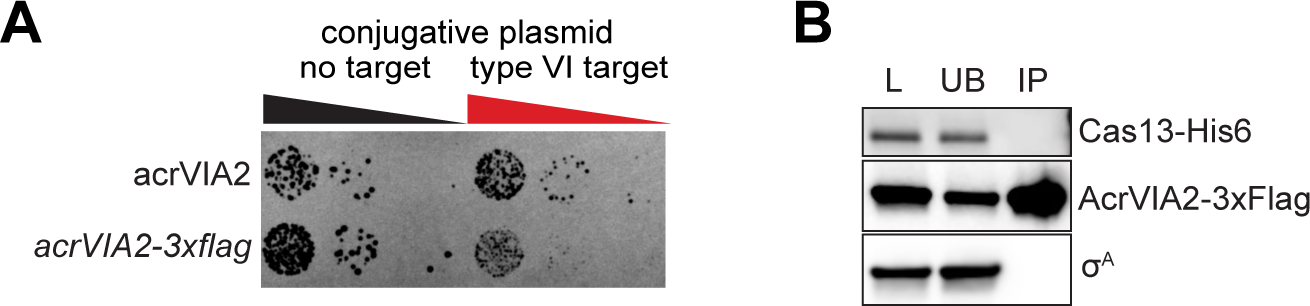
No detectable interaction between AcrVIA2 and Cas13. **(A)** AcrVIA2-3xFlag is partially functional in inhibition of immunity against a plasmid expressing an RNA protospacer recognized by the type VI-A CRISPR system. **(B)** No detectable co-immunoprecipitation of Cas13-his6 and AcrVIA2-3xFlag. The housekeeping sigma factor cr^A^ is shown as a non-interacting control. L, load, UB, unbound, IP, immunoprecipitate.

**Figure S15.**
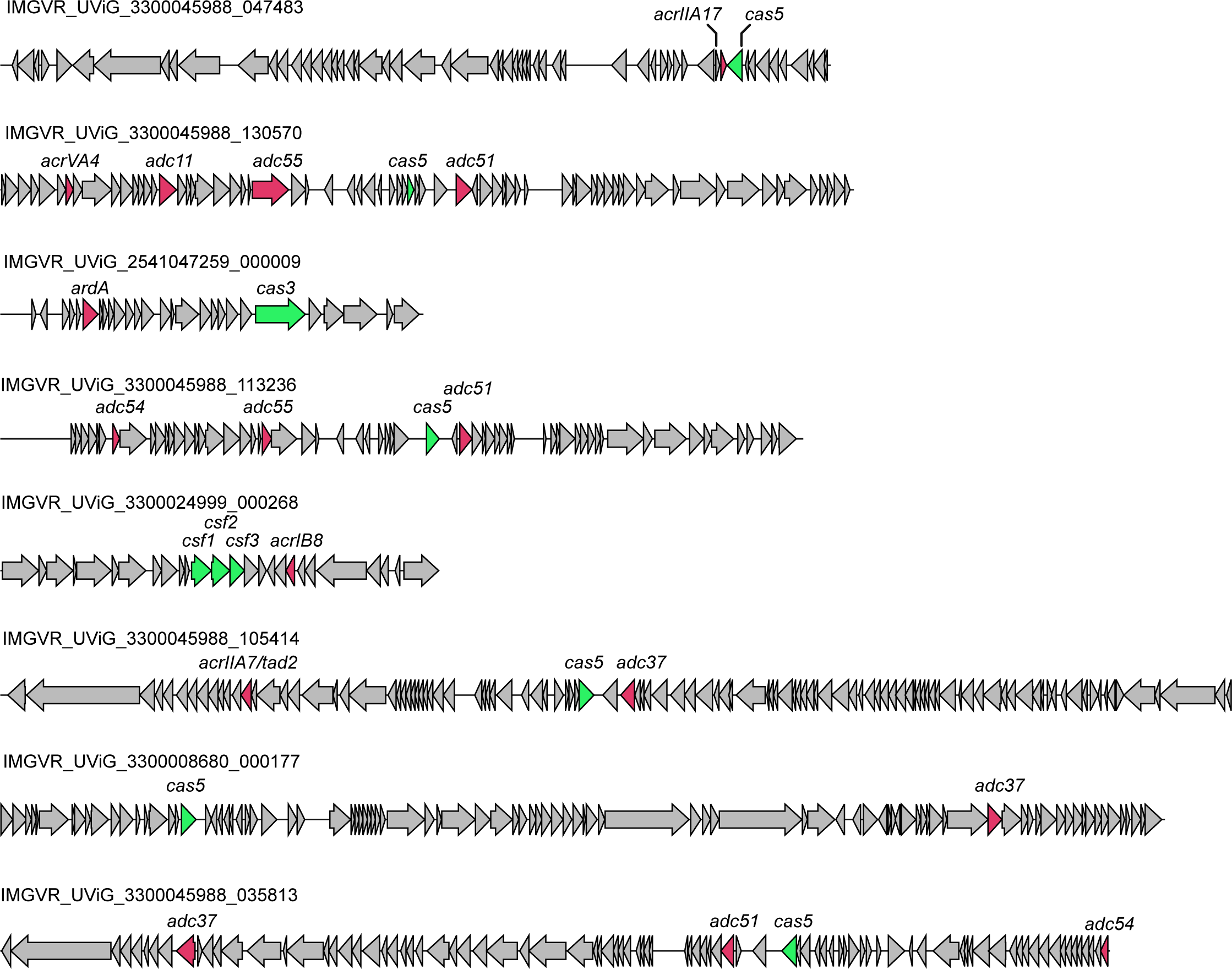
Orphan viral cas genes in the vicinity of known *acr* genes or putative anti-defense candidates (*adc*). Orphan viral cas gene homologs detected in the IMGVR database (green) that are located nearby known anti-CRISPR genes or putative anti-defense candidates (red). The unique viral genome identifer (UViG) number is shown for each locus. See other examples in Fig. 5B.

